# Multiscale mechanics drive functional maturation of the vertebrate heart

**DOI:** 10.1101/2024.07.24.604962

**Authors:** Toby G R Andrews, Jake Cornwall-Scoones, Marie-Christine Ramel, Kirti Gupta, James Briscoe, Rashmi Priya

## Abstract

How simple tissue primordia sculpt complex functional organs, robustly and reproducibly, remains elusive. As the zebrafish embryo grows into a larva, to improve its heart function, the embryonic myocardial wall transforms into an intricate 3D architecture, composed of an outer compact layer enveloping an inner layer of multicellular trabecular ridges. How these tissue layers acquire their characteristic form suited for their function remains an open question. Here, we find that multiscale mechanochemical coupling and an emergent tissue-scale morphological transition steer functional maturation of the developing zebrafish heart. Single-celled trabecular seeds recruit outer compact layer cells to mature into clonally heterogenous multicellular ridges, thereby amplifying cardiac contractile forces. In response, remaining compact layer cells are stretched, which impedes their further recruitment, thereby constraining trabecular ridge density. Concomitantly, Notch-dependent actomyosin dampening triggers a sharp transition in myocardial tissue area, activating rapid organ growth that expands blood filling capacity. Thus, multiscale self-organizing interactions optimize heart size and contractile efficiency to support embryonic life.

## Introduction

A hallmark of embryonic development is the formation of organs with reproducible shape, size, and function, essential for organismal growth and life. Yet, our understanding of how developing organs build complex anatomical structures to support their function remains limited. This is partly because previous studies have largely focussed on the genetic basis of organ development or studied tissues with relatively simple and static geometries^1–7^. These powerful approaches have uncovered key morphogenetic principles, but they do not fully recapitulate the complex dynamic nature of visceral organ morphogenesis^5,8–19^. The developing zebrafish heart is an exemplary model system to ask how organ form and function emerge during embryogenesis. Its tractability for imaging, perturbations and functional measurements enables a holistic understanding of single cell behaviours in the context of organ form and function^10,20–23^.

As the zebrafish embryo grows into a larva, to sustain its increasing physiological demands, the cardiac chambers undergo morphogenetic maturation to improve heart function^5,24^. During chamber maturation, the ventricular myocardial wall transforms from a simple epithelial monolayer into a complex 3D architecture composed of two distinct layers: an outer compact layer enveloping an inner trabecular layer **(Fig. 1A-A**’) ^5,24–27^. We previously showed that local differences in actomyosin tension trigger stochastic delamination of single cells from the outer compact layer to seed the inner trabecular layer^25^. As development proceeds, single trabecular seeds focally grow in cell number, elongate across the myocardial wall and project into the ventricle lumen, collectively building a 3D topological meshwork **(Fig. 1A-A**’) ^5,24,25^. Optimal density and organization of trabecular ridges is crucial for heart function^24,26,28,29,30^. Yet, how single trabecular seeds build multicellular ridges remains poorly understood. Meanwhile, the outer compact layer is a continuous cellular monolayer that defines ventricle size and shape and sustains extreme mechanical deformations during each heartbeat^31^. Despite its structural and functional importance, morphogenesis of the compact layer has been largely neglected. Thus, how myocardial morphogenesis is orchestrated to build a functional beating heart remains an open question.

**Figure 1.**
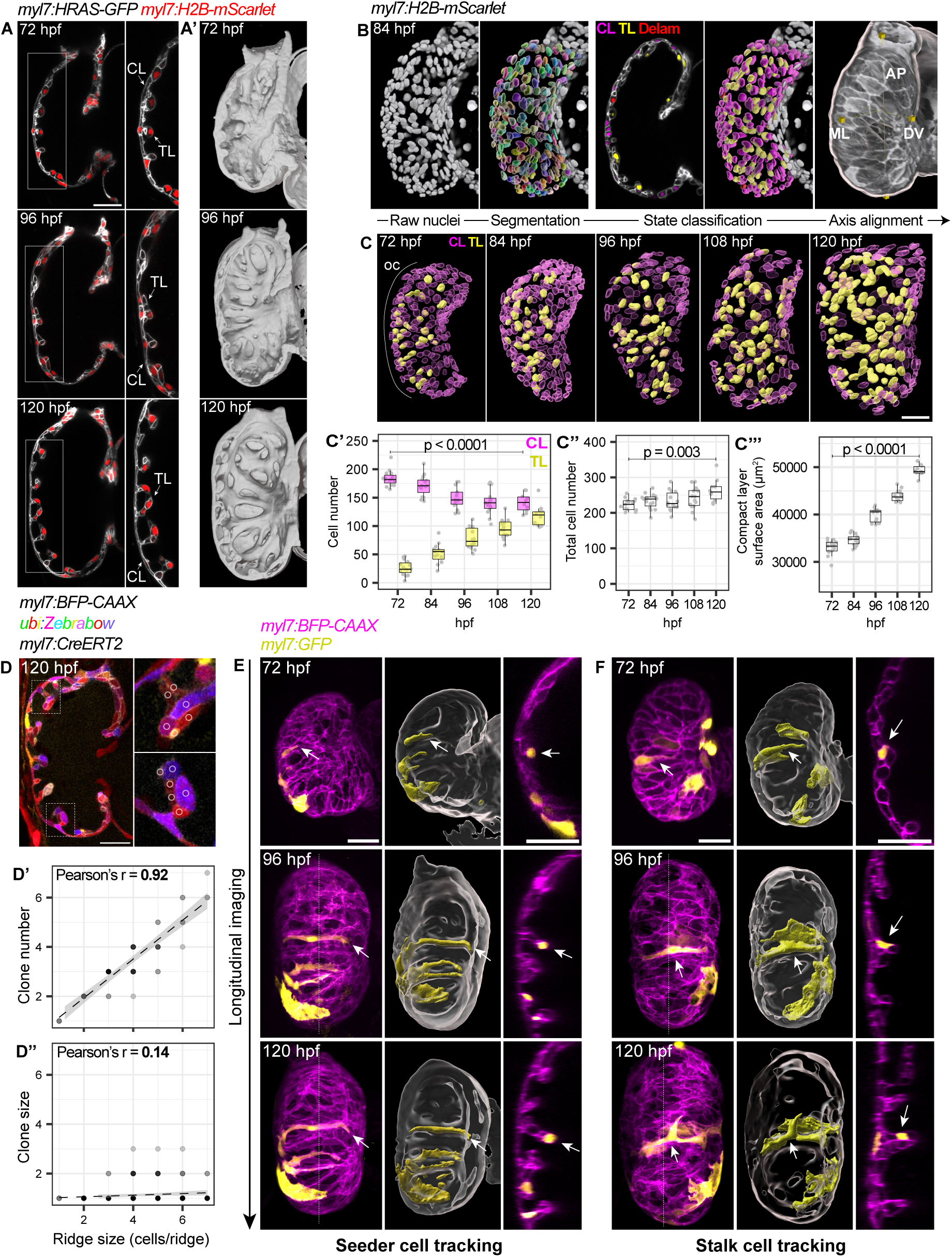
Compact layer cell recruitment dominates over proliferation to drive trabecular ridge growth. (A-A’) Midsagittal sections (A) and 3D surfaces (A’) of *myl7:HRAS-GFP* (membrane); *myl7:H2B-mScarlet* (nuclei) ventricles showing maturation of the trabecular (TL) and compact layer (CL), marked with arrows. (B) Image processing pipeline to quantify number and spatial organisation of cardiomyocyte nuclei (compact layer, CL; trabecular layer, TL; delaminating, delam; anteroposterior, AP; mediolateral, ML; dorsoventral, DV). (C-C”’) Nuclear segmentation time series (C), with quantification of CL and TL cell number (C’), total cell number (C”), and CL surface area (C”’). OC, outer curvature. n = 19,569 cells, 15 embryos per stage. P values result from unpaired two-tailed Students’ t-tests. (D-D”) Midsagittal section of *myl7:BFP-CAAX* (membrane); *ubi:Zebrabow-M* ventricle showing clonal heterogeneity in trabecular ridges after recombination with 10 µM 4-OHT from 60 - 72 hpf (inlays show ROIs for colour measurement), with correlation of ridge size (cells/ridge) with number of unique clones (D’) and clone size (cells/clone) (D’’). n = 247 cells, 64 ridges. (E, F) Maximum intensity projections (MIPs), 3D renderings and sections (midsagittal, or orthogonal to dashed lines) of ventricles expressing mosaic GFP. Representative TL seeds (E, representative of n=10/13) and stalk cells (F, representative of n=26/32) tracked longitudinally. White arrows mark tracked cells. Scale bars = 30 µm.

Here, we used 4D *in toto* live imaging, multiscale morphometrics, predictive theoretical modelling and spatiotemporally controlled perturbations to reveal how the functional architecture of the zebrafish heart emerges during development. Single-celled trabecular seeds build clonally heterogenous multicellular ridges by recruiting cells from the outer compact layer, thereby increasing myocardial muscle mass and contractile efficiency. As cardiac contractile forces increase, compact layer cells are irreversibly stretched, which increases myocardial tissue surface area in the absence of significant proliferation. In an autoinhibitory loop, cell stretching inhibits their further recruitment for ridge growth, thereby self-organising trabecular ridges to a set point density. Combining theory and experiments, we find that Notch mediated local actomyosin dampening triggers a sudden increase in myocardial tissue surface area. Once a threshold fraction of cells have activated Notch signalling, global tissue tension drops to a tipping point, at which cells collectively yield to cardiac forces and stretch. Of note, this non-linearity arises from mechanical coupling between cells, which averages effective tension across the tissue. This transition results in rapid and stable organ-scale growth that expands blood filling capacity. Together, our findings show that multiscale synergistic interactions between mechanics, signalling and organ function drives morphological and functional maturation of the developing vertebrate heart.

## Results

### Compact layer cell recruitment dominates over proliferation to drive trabecular ridge growth

Between 72 and 120 hpf, ventricle chamber maturation is characterized by increase in chamber size and growth of trabecular ridges, that elongate across the chamber wall and project into the lumen **(Fig. 1A-A’)**. To understand how these two morphogenetic processes are coordinated over time, we began with a morphometric analysis of myocardial tissue architecture. 3D coordinates of cardiomyocyte nuclei were mapped in an atlas of ventricles imaged every 12 hours between 72 and 120 hpf (hours post-fertilization), enabling quantitative analysis of changes in ventricle shape, size and cellular spatial organisation **(Fig. 1B, C, S1A)**. At 72 hpf, trabecular seeds were enriched in the outer curvature, as previously reported **(Fig. 1C, S1B, B’)** ^25,32^. In the following 48 hours, as single trabecular seeds transformed into multicellular ridges and dispersed across the ventricle wall **(Fig. S1B’)**, trabecular cell number increased by 4.5-fold **(Fig. 1C’)**. Strikingly, growth in trabecular cell number coincided with a 27% decrease in the number of outer compact layer cells **(Fig. 1C’)**, meaning total cell number increased by only 17% **(Fig. 1C’’)**. Despite losing cells, compact layer surface area expanded 51% during trabecular growth **(Fig. 1C’’’)**, enabling a 92% inflation in total ventricle volume **(Fig. S1C-C’)**.

To resolve which cellular processes transform single trabecular seeds into multicellular ridges, we first focussed on clonal dynamics using Brainbow technology (*Zebrabow-M*)^33^. We observed conspicuous clonal heterogeneity in trabecular ridges as early as 120 hpf **(Fig. 1D)**. Using a quantitative approach to classify clonal IDs **(Fig. S1D)**, we found that ridge size correlates strongly with the number of unique clones in each ridge **(Fig. 1D’)**, but only weakly with clone size **(Fig. 1D’’)**. These results implied a minor role for cell division in trabecular growth. We therefore focussed on local cellular rearrangement by mosaically labelling and tracking trabecular cells. In agreement with our clonal analysis **(Fig. 1D-D’’)**, GFP-positive trabecular seeder cells did not divide to form GFP-positive ridges, and instead persisted at the tips of maturing ridges composed of GFP-negative cells **(Fig. 1E)**. Retrospective tracing of stalk cells showed that they originate from the compact layer and enter the ridge base as it elongates and matures **(Fig. 1F)**. We further confirmed these observations by tracking single trabecular nuclei during ridge growth using the photoconvertible protein KikGR^34^ **(Fig. S1E)**. Photoconverted trabecular seeds persisted at the tips of maturing ridges and rarely divided **(Fig. S1E, E’)**. We therefore conclude that trabecular seeds build clonally heterogenous ridges through local cellular rearrangement, in which compact layer cells iteratively enter the bases of the maturing ridges. This reasoning is consistent with our observation that the compact layer loses cells during trabecular growth **(Fig. 1C’)**. However, it also poses an open question of how the compact layer sustains tissue-scale growth and expands its surface area by 51% **(Fig. 1C”’)**, while losing a significant fraction of its cells.

### Trabecular ridge maturation triggers stretching of compact layer cells

To understand how compact layer area increases despite losing cells to fuel trabecular maturation, we needed a deeper understanding of its cellular morphology. Due to the inherent curvature of the ventricle surface, analysing the properties of compact layer cells has remained challenging. Accordingly, its properties have been neglected, or inferred indirectly from 2D sections of the ventricle^35^. Thus, we established a bespoke image processing workflow to isolate the ventricle surface, flatten it into a pseudo-2D sheet, segment individual compact layer cells and measure their geometric features **(Fig. 2A, S2A, B)**. Using this quantitative approach, we found that despite striking diversity in their morphology, the mean area of compact layer cells increased 69% during trabecular maturation **(Fig. 2B, B’)**. To understand if this area expansion results from volumetric cell growth, we tracked changes in the cytoplasmic volumes of single compact layer cells and measured a slight increase (8%) **(Fig. 2C, C’)**, which cannot account for the 69% increase in surface area between 72 and 120 hpf **(Fig. 2B’)**. We therefore reasoned that compact layer cells should expand their apical area by changing shape. Indeed, expansion of cell area coincides with a 76% decrease in apicobasal thickness, reinforcing that cells mainly expand their area by stretching, while preserving their volume **(Fig. 2D, D’)**.

**Figure 2.**
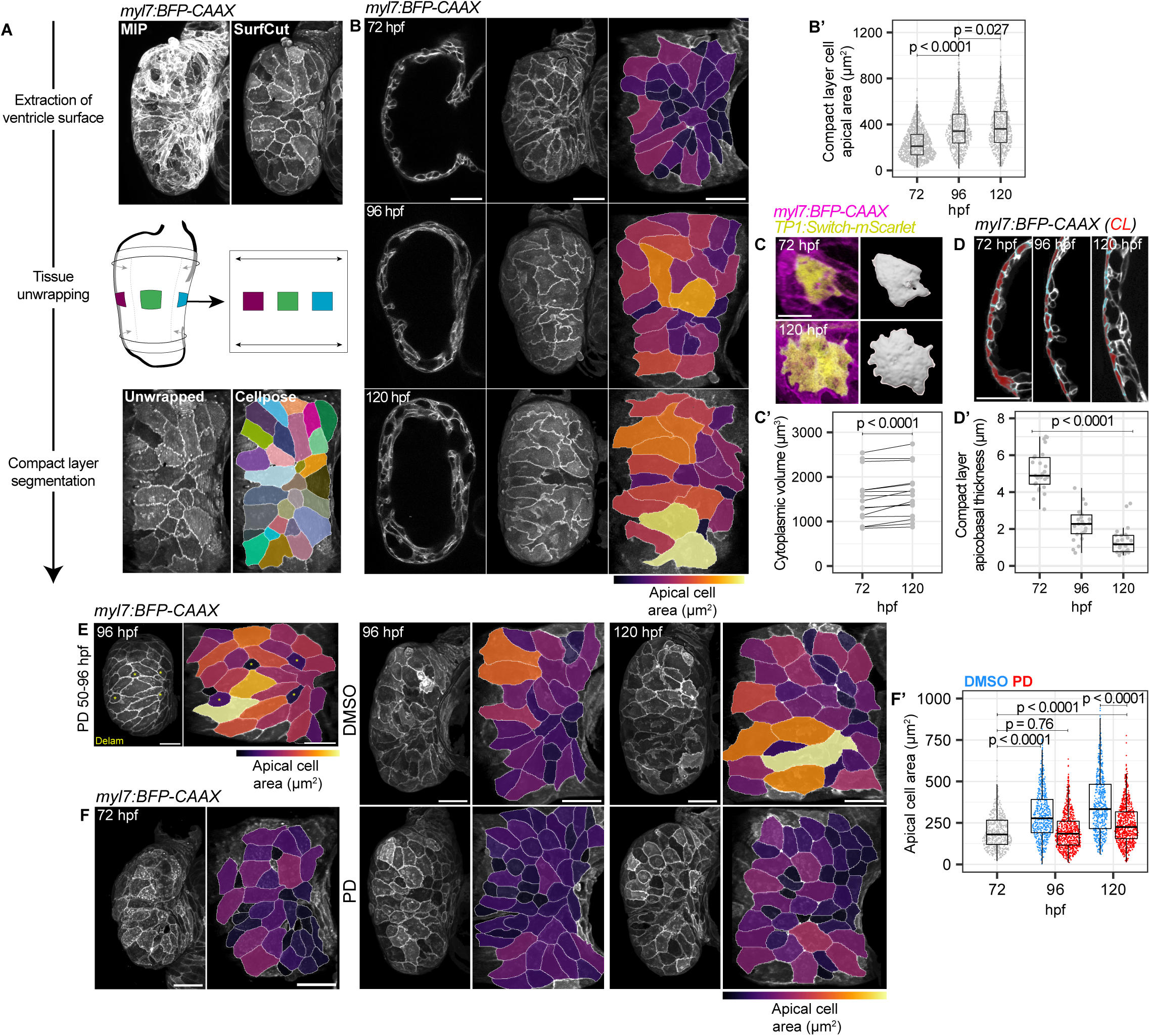
Trabecular ridge maturation triggers stretching of compact layer cells. (A) Summary of image processing pipeline for geometric quantification of CL cells: isolation of CL signal, unwrapping to reduce geometric distortion, segmentation of cell apical domains, quantification of geometric features. (B-B’) Midsagittal sections, CL surface and flat projections of *myl7:BFP-CAAX* (membrane) ventricles with cell area heatmaps (B), and area quantification (B’). n = 2922 cells. P values result from unpaired Mann-Whitney U tests. (C-C’) Longitudinal imaging of CL cells using mosaic *TP1:Switch-mScarlet* expression (C), with volume quantification per cell (C’) n = 15 cells. P value results from unpaired two-tailed Students’ t-test. Scale bar = 20 µm. (D-D’) Midsagittal sections of outer curvature of 72 – 120 hpf ventricles, with compact layer cells pseudocolored (D), and quantification of CL thickness (D’). n = 25 measurements per stage. P value results from unpaired two-tailed Students’ t-test. Scale bar = 20 µm. (E) CL surface and flat projection with cell area heatmap for embryo treated with 10 µM PD168393 (PD, Erbb2 inhibitor) from 50-96 hpf. Yellow asterisks mark delaminating cells (Delam). Representative of n = 6 embryos. (F-F’) CL surface and flat projections with cell area heatmaps for ventricles treated with 10µM PD168393 (PD) (F), quantification of CL cell area (F’). 72 hpf, n = 18 embryos; 96 hpf DMSO, n = 20 embryos; 120 hpf DMSO, n = 21 embryos; 96 hpf PD, n = 21 embryos; 120 hpf PD, n = 21 embryos. P values result from unpaired Mann-Whitney U tests. Scale bars = 30 µm, unless otherwise stated.

We next asked what drives compact layer cell stretching. A careful analysis of the compact layer surface at 84 hpf revealed that cells with large apical areas exhibit a rosette-like organisation around delaminating cells with reduced apical area **(Fig. S2C)**, which seed trabecular ridges^25^. Moreover, using a chemical ErbB2 inhibitor to restrict delamination to only 1-5 cells on a naïve myocardium, we found stretched cells are concentrated around delaminating cells **(Fig. 2E)**. These observations indicated a spatial correlation between delamination and compact layer cell stretch. Next, to test whether compact layer stretch is functionally coupled to trabecular maturation, we treated embryos with a chemical ErbB2 inhibitor to block trabecular maturation, as shown previously^36^, and analysed its effect on compact layer morphology at successive stages of stretch **(Fig. 2F)**. Compared to DMSO-treated hearts, ErbB2 inhibited ventricles retained small, rounded compact layer cells up to 120 hpf, characteristic of untreated ventricles at 72 hpf **(Fig. 2F)**. Using our quantitative pipeline **(Fig. S2A)**, we found that blocking trabeculation reduces compact layer cell stretch by 57% **(Fig. 2F’)**. Similarly, cells also failed to stretch in trabeculation-deficient *nrg2a* mutant embryos, that lack the upstream ligand for ErbB2 signalling **(Fig. S2D, D’)** ^32^. These results implied that, as well as depleting cells from the compact layer as they form, maturation of trabecular ridges cause remaining compact layer cells to stretch. We then sought to understand how these latter steps were linked.

### Multiscale feedback drives compact layer remodelling and constrains trabecular density

Trabeculation has been postulated to increase cardiac contractile force production and blood flow ^5,28^. We therefore reasoned that trabecular maturation might also promote compact layer cell stretching by increasing global cardiac forces. Given that a direct quantitative correlation between trabecular maturation and cardiac function is still missing, we first performed a systematic measurement of changes in cardiac physiology during trabecular maturation using timelapse imaging of beating hearts **(Fig. 3A, Supplementary Video 1)**. Between 72 and 120 hpf, as trabecular ridges mature, stroke volume increased by 36% **(Fig. 3A’)**, resulting from a 50% increase in end-diastolic volume **(Fig. 3A’’)** and 28% increase in ejection fraction **(Fig. 3A’’’)**. Of note, this functional maturation was abrogated when trabecular growth was blocked by ErbB2 inhibition **(Fig. 3A’-A’’’, Supplementary Video 1)**. At 120 hpf, stroke volume was equal to that measured at 72 hpf **(Fig. 3A’)**, while ejection fraction deteriorated significantly in the absence of trabecular maturation **(Fig. 3A’’’)**. Overall, these observations indicate that trabecular maturation improves the functional efficiency of the ventricle by increasing its filling capacity and contractile forces.

**Figure 3.**
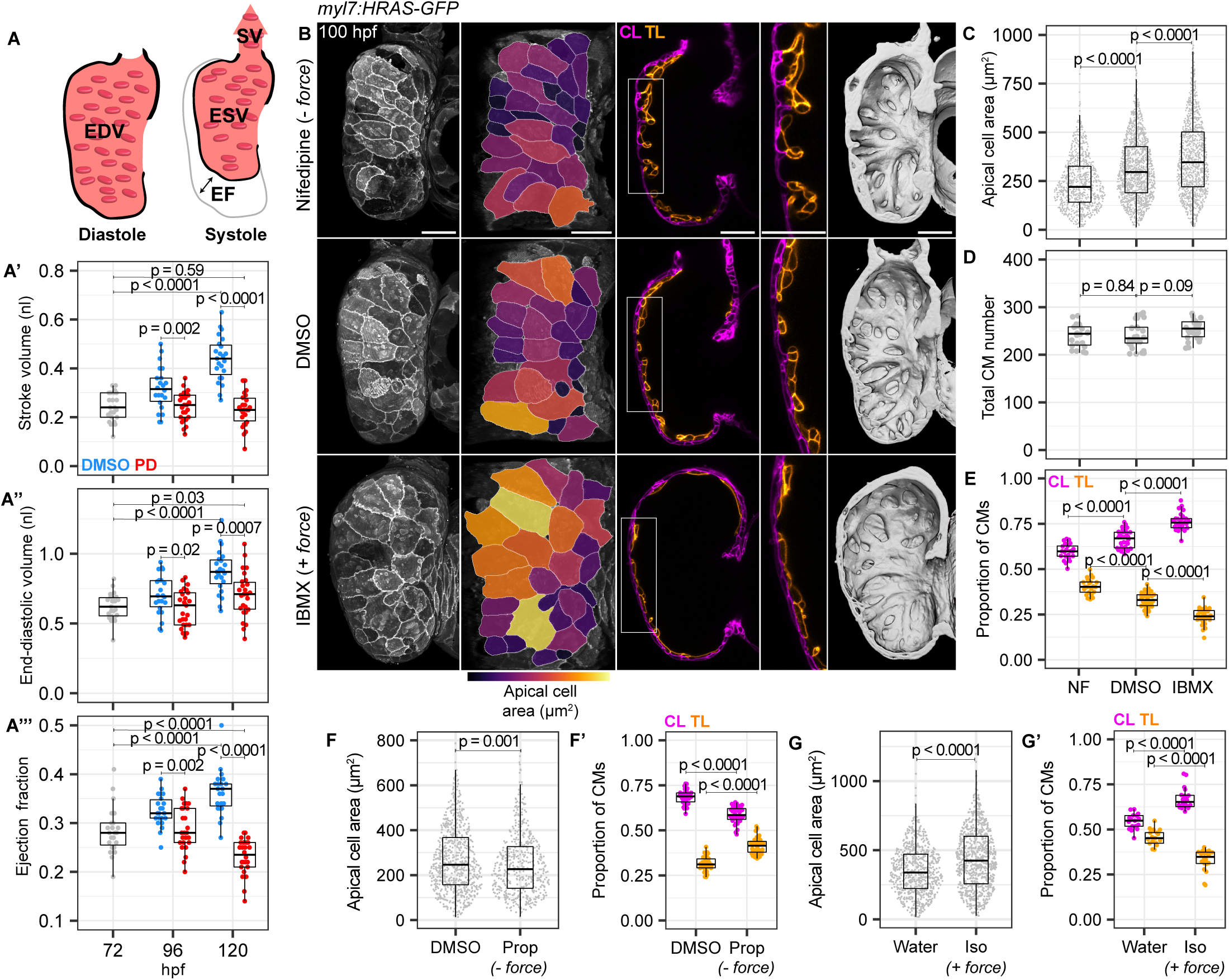
Multiscale feedback drives compact layer remodelling and constrains trabecular density. (A-A’’’) Schematic for inferred cardiac functional parameters (A), end-diastolic volume, EDV reflects blood filling capacity; end-systolic volume, ESV; ejection fraction, EF is a scale-free measure of contractile efficiency; stroke volume, SV is a function of ventricle filling capacity and contractile efficiency. Quantifications for embryos treated with DMSO or PD168393 (PD) (A-A’’’). n = 23 embryos, 72 hpf; 22 embryos, 96 hpf DMSO; 23 embryos, 120 hpf DMSO; 25 embryos, 96 hpf PD; 26 embryos, 120 hpf PD. P values result from unpaired Students’ t-tests. (B-E) CL surface, flat projections with cell area heatmaps, pseudocoloured midsagittal sections, cropped sections (white boxes), and 3D renderings of *myl7:HRAS-GFP* (membrane) ventricles treated with 10 µM Nifedipine (NF), DMSO or 50 µM IBMX from 80-100 hpf (B), with quantification of cell area (C), total cardiomyocyte (CM) number (D) and proportion of CL and TL CMs (E). n = 30 embryos, DMSO; 22 embryos, NF; 26 embryos, IBMX. P values result from unpaired Students’ t-tests. (F-F’) Quantification of cell area (F) and proportion of CL and TL CMs (F’) for embryos treated with DMSO or 100 µM Propranolol (Prop) from 80-100 hpf. n = 30 embryos, DMSO; 34 embryos, Prop. P values show result of unpaired Mann-Whitney U tests (F) and unpaired Students’ t-tests (F’). (G-G’) Quantification of cell area (G) and proportion of CL and TL CMs (G’) for embryos treated with water or 50 µM Isoprenaline (Iso) from 80-100 hpf. N = 17 embryos, water; 20 embryos, Iso. P values show result of unpaired Mann-Whitney U tests in (G) and unpaired Students’ t-tests in (G’). Scale bars = 30 µm.

We then asked whether changes in global cardiac force production influence stretching of compact layer cells. To this end, we modulated cardiac force production using orthogonal pharmacological approaches: Nifedipine (Ca^2+^ channel blocker) and Propranolol (β-receptor antagonist) acutely decreased stroke volume ^37,38^, whereas IBMX (phosphodiesterase inhibitor) and Isoprenaline (β-receptor agonist) acutely increased stroke volume **(Fig. S3A, Supplementary Video 2-3)** ^39^. Reducing cardiac forces decreased compact layer cell area and increased apicobasal thickness **(Fig. 3B, C, F, S3B, D).** Conversely, increasing cardiac force increased compact layer cell area and decreased apicobasal thickness **(Fig. 3B, C, G, S3C, D)**. Together, these results indicate that compact layer cells stretch in response to increase in global organ-scale force production induced by trabecular growth **(Fig. 3B-C),** as well as local cellular rearrangements driving trabecular maturation **(Fig. 2E-F’).**

Surprisingly, manipulating cardiac force had an inverse effect on trabecular growth. While increasing cardiac force accelerated compact layer stretching **(Fig. 3B,C)**, trabecular ridge complexity appeared reduced; ridges were smaller and flattened against the chamber wall **(Fig. 3B, S3C)**. Conversely, decreasing cardiac force abrogated compact layer stretching, but increased trabecular complexity, and ridges projected further into the ventricle lumen **(Fig. 3B, S3B)**. Of note, total cell number remained unchanged in these manipulations, suggesting that the effect on trabecular complexity was not a result of changes in proliferation **(Fig. 3D, S3B’, C’)**. Instead, spatial allocation of cells between the compact and trabecular layers was disrupted, where stretching of compact layer cells abrogates their further recruitment for trabecular growth **(Fig. 3E, F’, G’)**.

Collectively, these findings reveal multiscale mechanical feedback regulating compact and trabecular layer maturation. Recruitment of compact layer cells drives trabecular maturation, resulting in local cell shape changes and an increase in global cardiac force production. In response, remaining compact layer cells are stretched, which impedes their recruitment for trabecular growth. This negative feedback loop stabilizes trabecular density to a physiological set point important for heart function, while promoting cell shape changes associated with ventricle growth.

### Stretched cells deplete their junctional actomyosin pool to enable myocardial tissue growth

As cell shape is influenced by both extrinsic forces and intrinsic mechanical properties^40,41^, we next asked whether compact layer cell stretching is actively regulated by actomyosin mechanics. To this end, we generated a myocardial Lifeact-mNeongreen reporter line to analyse actin organisation in live compact layer cells and used phalloidin to label the endogenous actin pool. At 72 hpf, actin was strongly enriched at cell-cell junctions in a purse string organisation **(Fig. 4A)**. Using a quantitative image analysis pipeline **(Fig. S4A)**, we find that as cells stretch between 72 and 120 hpf, junctional actin was depleted, and a relatively homogenous actin network emerged across the cell **(Fig. 4A-A’, S4B-B’)**. By comparing the area and actin organisation of single compact layer cells, we found that junctional actin enrichment and area expansion were mutually exclusive properties, suggesting cells may first deplete their junctional actin before expanding in area **(Fig. S4D)**. Of note, this change in actin organisation did not result from its global depletion or changes in its sarcomeric structure **(Fig. S4B’’, C)**. Similar to actin, we also observed junctional depletion of the sarcomeric Z-disk scaffold protein actinin3b (actn3b) in stretched cells **(Fig. 4B, B’)**. Of note, analysis of actn3b organisation revealed a progressive uniform alignment of sarcomeres to the mediolateral axis of the ventricle **(Fig. S4E-E’)**. This alignment mirrored the primary axis of ventricle dilation during diastole, suggesting that reorientation of sarcomeres might support stronger contraction along this axis for efficient ejection of blood **(Fig. S4 E’’)**.

**Figure 4.**
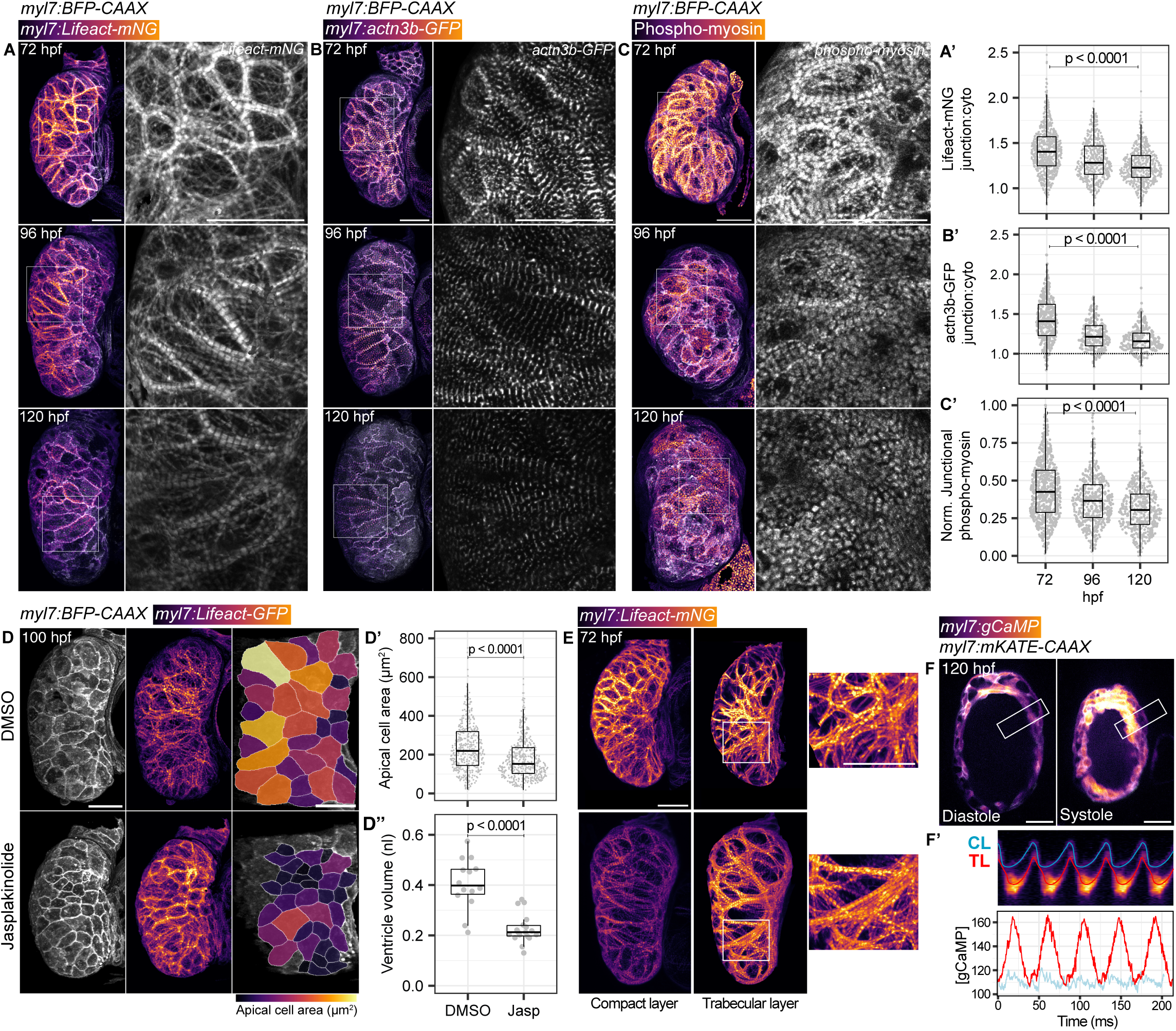
Stretched cells deplete their junctional actomyosin pool to enable myocardial tissue growth. (A-C’) CL surface in *myl7:BFP-CAAX* (membrane) ventricles, showing intensity (in heatmap) and organization (greyscale cropped panels) of actin (Lifeact-mNG) (A), actinin (actn3b-GFP) (B) and phospho-myosin (C), with quantifications of junction:cytoplasmic ratio (A’, B’), or normalised junctional fluoresence (C’). Lifeact-mNG: 72 hpf, n = 757 cells; 96 hpf = 521 cells; 120 hpf = 541 cells. actn3b-GFP: 72 hpf, n = 343 cells; 96 hpf, 204 cells; 120 hpf, 214 cells. Phospho-myosin: 72 hpf, n = 802 cells; 96 hpf = 498 cells; 120 hpf = 479 cells. P values result from unpaired Mann-Whitney U tests. (D-D”) CL surface and flat projections with cell area heatmap for ventricles treated with DMSO or 0.5 µM Jasplakinolide (Jasp) from 80-100 hpf, with quantification of Lifeact-GFP junction:cytoplasm ratio (D’) and CL cell area (D’’). n = 412 cells, DMSO; n = 511 cells, Jasp. P values result from unpaired Mann-Whitney U test (D’) and unpaired Students’ t-test (D’’). (E) Maximum intensity projections and manually segmented actin (Lifeact-mNG) in the compact and trabecular layer. Heatmaps scaled per column. Cropped panels (white boxes) show actin organization in trabecular ridges at each stage. (F-F’) Stills from beating heart movie at 120 hpf, with kymograph (F’) through white box in (F), and line quantification for changes in gCAMP fluoresence. Representative of n = 11 hearts. Scale bars = 30 µm.

Since junctional actomyosin organisation influences junctional tension^42^, we analysed two canonical markers of junctional tension; phosphorylated myosin light chain (phospho-myosin), which labels the active myosin pool **(Fig. 4C)**, and vinculin, which is recruited to cadherin junctions in a load-dependent manner **(Fig. S4F)**^43^. We measured a decline in junctional phospho-myosin intensity in compact layer cells between 72 and 120 hpf **(Fig. 4C’)**. In parallel, we developed a transgenic reporter line to localise vinculin, and found junctional vinculin decreases as compact layer cells stretch **(Fig. S4F-F’)**. Together, we conclude that stretched compact layer cells dilute their junctional actomyosin and lower their junctional tension. We therefore reasoned that actomyosin remodelling may permit cells to stretch. To test this experimentally, we globally inhibited actin turnover by treating embryos with jasplakinolide **(Fig. 4D)**. Jasplakinolide treatment from 80 to 100 hpf stabilized the junctional actin purse string in compact layer cells **(Fig. 4D, S4G)**, and abrogated cell area growth by 37% **(Fig. 4D’)**. Strikingly, jasplakinolide treatment also blocked whole ventricle growth, reflected by a 47% lower volume than untreated embryos **(Fig. 4D’’)**. Collectively, these results suggest that junctional actomyosin depletion in compact layer cells enables them to stretch and expand myocardial tissue area.

Importantly, the compact and trabecular layers exhibit distinct dynamics of actin remodelling **(Fig. 4E)**. While compact layer cells dissolve their junctional actin, trabecular cells assemble dense linear sarcomeric actin bundles oriented along the length of each ridge **(Fig. 4E)**. We hypothesized that differential actomyosin organization could drive an emergent functional distinction between each cardiomyocyte layer – as the compact layer lowers its tension and stretches, the trabecular layer adopts the primary contractile function. Since the amplitude of Ca^2+^ transients in cardiomyocytes determines the strength of contraction ^44^, we analysed their dynamics using live imaging of a genetically encoded GCaMP biosensor **(Fig. 4F, Supplementary Video 4)**. While both compact and trabecular layers show coherent Ca^2+^ transients at 120 hpf, the amplitude was much greater in the trabecular layer, indicating a stronger stimulus for contraction **(Fig. 4F’)**. Together, these results reinforce that, as the ventricle matures, each cardiomyocyte layer becomes morphologically and functionally specialized. While trabecular maturation increases the myocardial muscle mass to amplify its contractile efficiency, compact layer cells lower their junctional tension allowing cell-scale stretch and organ-scale growth.

### Notch mediated actomyosin remodelling triggers irreversible stretching of compact layer cells

We next asked what triggers actomyosin remodelling to enable compact layer stretch. Since cell shape changes can feed back to cytoskeletal organisation, we tested whether compact layer cell stretching dilutes the junctional actin pool, as previously suggested^42,45,46^. We therefore ectopically stretched the compact layer cells by driving an early increase in cardiac force production with IBMX. However, IBMX treatment enriched rather than depleted junctional actin, despite increasing cell area **(Fig. S5A-A’’)**. Thus, cell stretch alone is not sufficient to remodel actin.

We therefore focused on Notch signalling, which has been shown previously to get exclusively activated in compact layer cells in a stochastic manner in response to delamination and mediates lateral inhibition to block ectopic delamination^25,26^. At 96 hpf, Notch-positive cells have a relatively homogenous actin organisation compared to neighbouring Notch-negative cells with enriched junctional actin **(Fig. 5A-A’)**. Notably, junctions between Notch-positive cells also exhibited weaker vinculin enrichment than those between Notch-negative cells, indicating that they are under lower tension **(Fig. 5B-B’)**. Experimentally, treating embryos with a chemical Notch inhibitor stabilized junctional actin **(Fig. S5B-B’)**, abrogated stretching of compact layer cells **(Fig. S5B’’)**, and reduced ventricle volume by 33% **(Fig. S5B’’’)**. This phenotype was similar to that observed with jasplakinolide treatment **(Fig. 4D-D”)**, further reinforcing that junctional actin depletion is required for cells to stretch and increase myocardial surface area. Moreover, mosaic expression of dominant-negative Suppressor of Hairless (DN-SuH), which desensitizes cells to Notch signalling^26^, caused cells to retain small, rounded morphologies and strong junctional actin enrichment, whereas DN-SuH negative cells dissolved their junctional actin and stretched **(Fig. 5C, C’)**. Conversely, we drove constitutive Notch activity using mosaic over-expression of the Notch intracellular domain (NICD)^26^ **(Fig. 5D)**. At 72 hpf, NICD expressing cells showed depleted actin fluoresence, consistent with accelerated actin remodelling **(Fig. 5D-D’)** and buckled outwards from the normal plane of the compact layer **(Fig. 5D),** indicating they have less actomyosin tension and thus are more deformable. Comparing NICD positive cell area with an adjacent region with the same radius predicted a mean 2-fold stretch occurring locally in NICD positive cells **(Fig. 5D’’)**. Considered together, these data implicate Notch signalling as a molecular trigger for actomyosin remodelling in compact layer cells which drives whole-ventricle growth.

**Figure 5.**
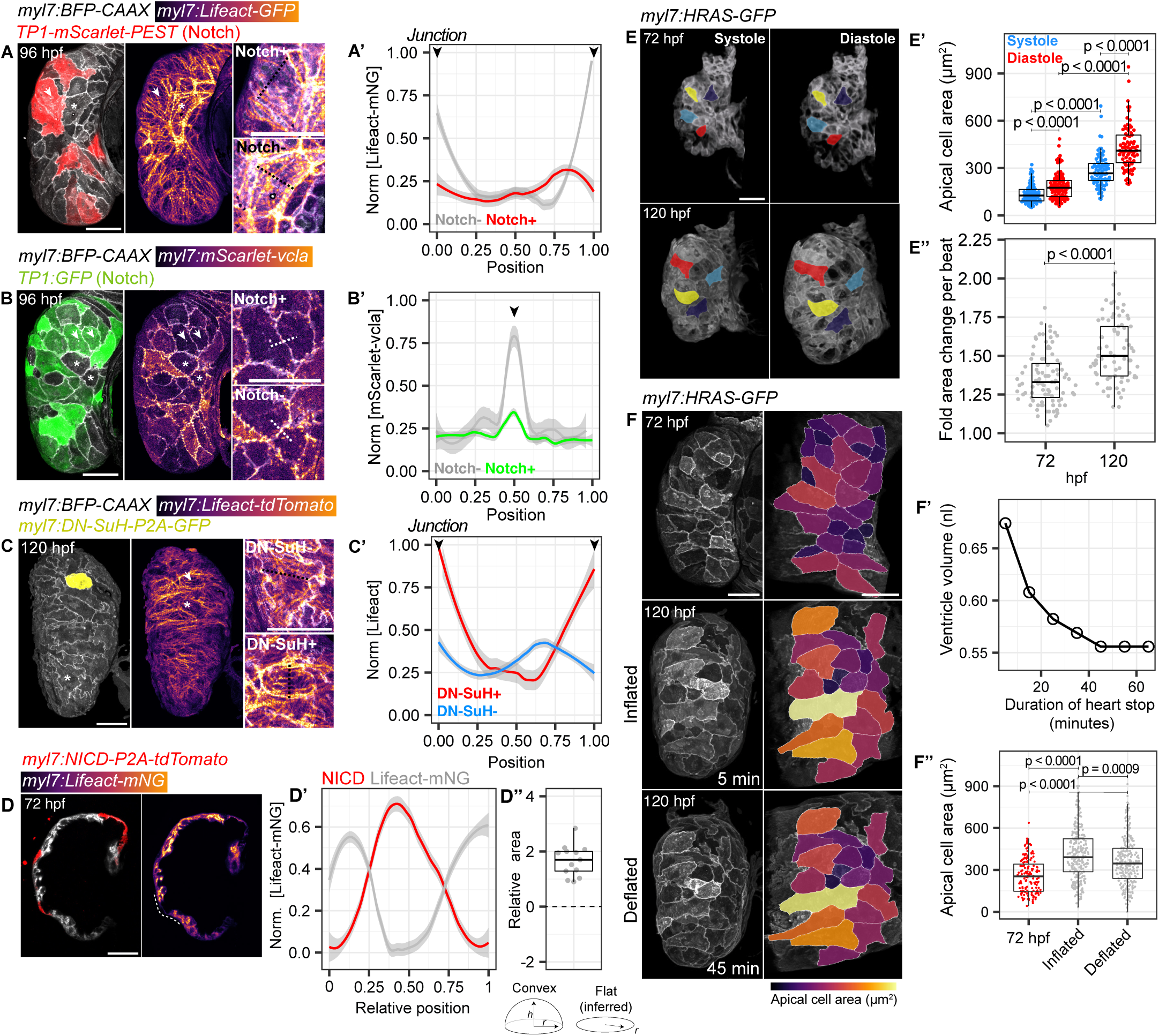
Notch mediated actomyosin dampening triggers irreversible stretching of compact layer cells. (A-A’) CL surface in *myl7:BFP-CAAX* (membrane) ventricles showing actin (Lifeact-GFP) in Notch+ve (arrow) and Notch-ve (asterisks) cells. Dashed lines mark line quantifications shown in (A’). Black arrowheads mark cell-cell junctions (A’). Representative of n = 17 ventricles. (B-B’) CL surface showing vinculin (mScarlet-vcla) localization at junctions of Notch+ve (arrow) and Notch-ve (asterisk) cells. Dashed lines mark line quantifications shown in (B’). Representative of n = 14 ventricles. (C-C’) CL surface with mosaic Notch inhibition (expression of DN-SuH). Inlays show actin (Lifeact-tdTomato) signal in DN-SuH-positive (arrow) and DN-SuH-negative (asterisk) cells. Dashed lines mark line quantifications shown in (C’). Representative of n = 12 ventricles. (D-D’’) Midsagittal sections of ventricles with mosaic NICD expression in cells bulging outwards from the CL (D), with quantification for actin (Lifeact-mNG) (D’) along dashed line in (D), and inference of relative cell area in NICD expressing domes compared to adjacent cells (D”). n = 13 measurements. (E-E”) CL surface from 3D beating heart movies in *myl7:HRAS-GFP* (membrane) ventricles at 72 and 120 hpf (sample cells tracked between phases pseudocoloured), with quantification of CL cell area in systole and diastole (E’), and fold cell area change between phases (E”). n = 93 cells, 72 hpf; n = 78 cells, 120 hpf. P values result from unpaired Mann-Whitney U tests. (F-F’’) CL surface and flat projections with cell area heatmap at 72 hpf, and 120 hpf +5 minutes or +45 minutes after heart-stop (F), with representative volumetric decrease in ventricle (F’), and quantification of CL cell area (F’’). n = 155 cells, 72 hpf; n = 278 cells, 120 hpf. P values result from unpaired Mann-Whitney U tests. Scale bars = 30 µm.

We next asked how molecular regulation of cell-scale mechanics impacts organ-scale material properties to facilitate ventricle growth. To this end, we acquired 3D movies of beating hearts at early (72 hpf) and late stages (120 hpf) of ventricle maturation **(Supplementary Video 5)**. At both stages, compact layer cell areas expanded to maximum values in diastole, and restored minimum values in systole **(Fig. 5E, E’)**, characteristic of reversible elastic deformations essential to resist short term perturbation and maintain cell shape^47^. Further, the extent of elastic deformation increased between 72-120 hpf **(Fig. 5E’’)**, congruent with cells depleting their junctional actomyosin, and an increase in ventricle volume and blood filling capacity **(Fig. 3A-A”’, 4A-C’)**. However, compact layer cells also cumulatively stretched on the order of hours to days **(Fig. 2B, B’)**, reflected by increased area throughout the heartbeat **(Fig. 5E’’)**. To test whether this long-term stretch represents an irreversible plastic deformation^47,48^, we relieved fluid pressure in the ventricle by stopping the heart and allowing it to collapse **(Fig. 5F, S5C)**. After 45 minutes, ventricles had fully collapsed **(Fig. 5F’)**, but mean compact layer cell area remained 39% greater than at 72 hpf, prior to the onset of stretch **(Fig. 5F’’)**, indicating an irreversible stretch. These results suggest that compact layer cells exhibit slow plastic deformations that yield irreversible tissue growth, a characteristic feature of developing tissues undergoing shape changes required for morphogenesis ^47,49^.

### Vertex model suggests Notch triggers sudden tissue growth through a global drop in tension

Our experimental data suggests that Notch mediated dampening of actomyosin tension allows compact layer cells to stretch in response to external force, resulting in organ-scale growth. Notch activation occurs mosaically in a small fraction of cells, leading to heterogeneity in cell shape and mechanics **(Fig. S6A, 2B, 5A)**^25^. In contrast, ventricle growth occurs rapidly **(Fig. 1C”’)**, and its shape and growth rate are highly reproducible between embryos **(Fig. S6B, B’)**. This raises an intriguing problem of how variable cellular dynamics produce stereotypical changes in organ-scale morphology^50–52^. To address this problem, we devised a minimal 3D vertex model, linking Notch induced dampening of junctional tension with tissue-scale growth dynamics **(Fig. 6A, A’**; see Supplementary Modelling section) ^53–55^. Here, we modelled the compact layer as a continuous sheet of cells exposed to a constant external stretching force **(Fig. 6A)**. In our model formulation, cells exhibit elastic behaviour below a threshold cell area, allowing them to resist compression^56^. Notch activation was then simulated by dampening junctional tension in a set frequency of cells (*pNotch*) **(Fig. 6A’’)**. By simulating our model to equilibrium, we tracked emergent changes in local cell geometry and global tissue growth dynamics ^57,58^.

**Figure 6.**
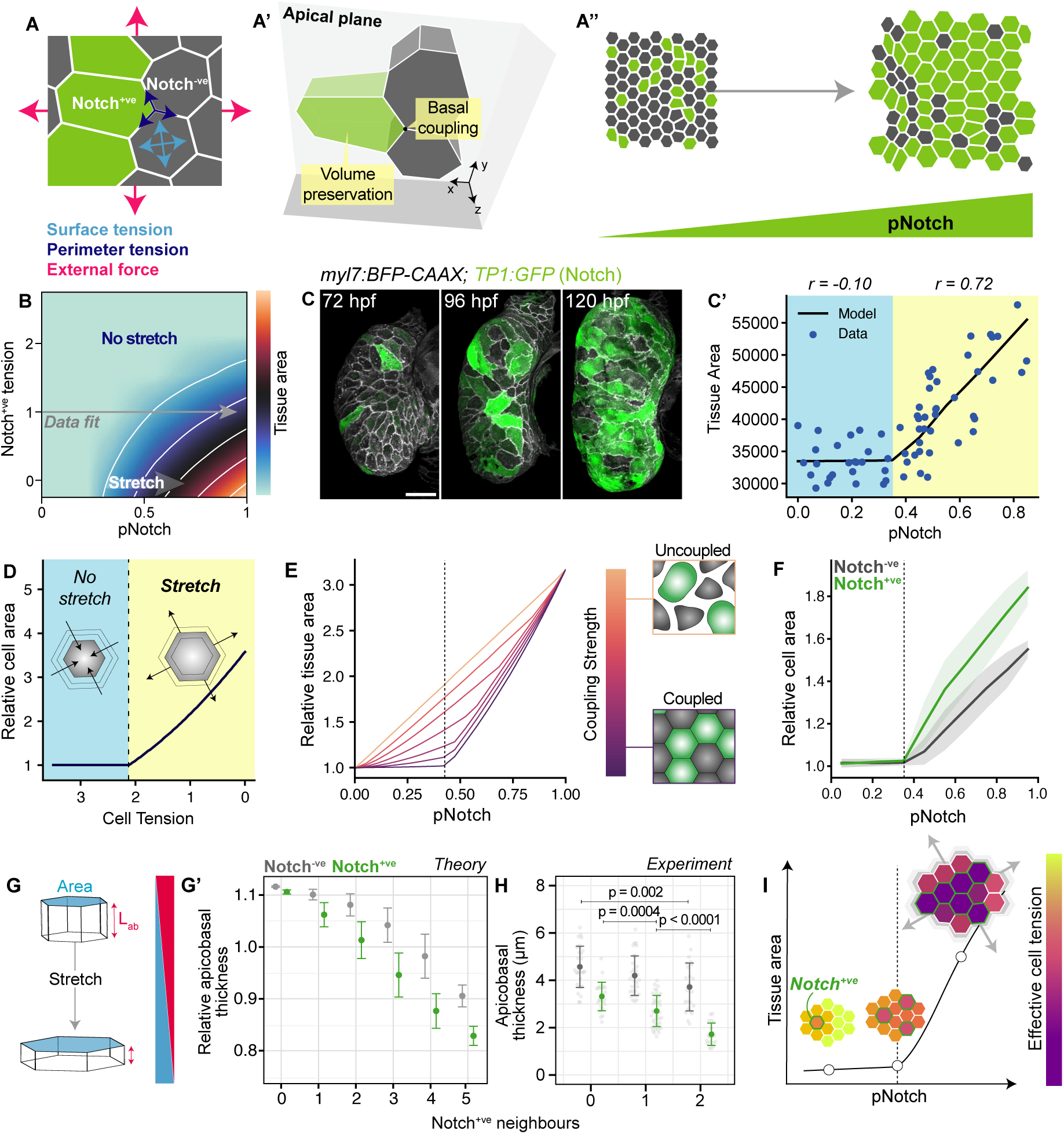
Vertex model suggests Notch triggers sudden tissue growth through a global drop in tension. (A-A’’) Schematic of model formulation showing constituent parameters: cell-intrinsic surface and perimeter tensions, and isotropic external force (A), with 3D representation (A’), and illustrated changes in pNotch (A”). (B) Subset of model phase space showing variation in tissue area (heatmap) with variation in the frequency (pNotch) and junctional tension of Notch+ve cells. White lines connect regions with same tissue area. Grey line shows closest fit of model to imaging data. (C-C’) CL projections of ventricles expressing *myl7:BFP-CAAX* (membrane) and a *TP1:GFP* Notch reporter, with quantification of compact layer cell area at different frequencies of Notch+ve cells (pNotch) and closest model fit (C’). n = 65 embryos. (D) Relationship between cell perimeter tension and area in cell-scale model, with variations in external force. Different tissue behaviour either side of tipping point (dashed black line) marked in blue (no stretch) and yellow (stretch) colour blocks. (E) Mean field simulation of model with different levels of cell coupling, showing relative tissue area at different levels of pNotch, with schematic for coupled and uncoupled states. (F) Quantification of relative cell area for Notch+ve and Notch-ve at increasing levels of pNotch in tissue-scale model. (G-H) Schematic showing inverse relationship between cell area and thickness (used as a readout of cell stretch where area is also influenced by differences in cell volume); Lab, cell apicobasal thickness (G), with quantification of changes in Notch+ve and Notch-ve cell thickness depending on the number of Notch+ve neighbours in tissue-scale model (G’, theory) and in vivo (H, experiment). in vivo: Notch-ve, n = 85 cells from 18 embryos; Notch+ve, n = 80 cells from 18 embryos. Error bars show mean and standard deviation. P values result from unpaired Students’ t-tests. (I) Schematic showing model of Notch-regulated tissue stretch. Stochastic Notch activity drives a global decrease in effective tissue tension (heat scale), to a tipping point where the system yields to cardiac forces and transitions from no stretch to tissue-wide stretch. Scale bars = 30 µm.

A 4D parameter sweep revealed multiple tissue stretch regimes governed by *pNotch* **(Fig. S6C)**. Depending on the balance of cell-intrinsic tensions and external forces, we observed variations in the rate and magnitude of tissue stretch in response to *pNotch* **(Fig. S6C).** Guided by our *in vivo* observations **(Fig. 3B, 5A-B’)**, we focused on a region of phase space where external force is strong and differential junctional tension between Notch^+ve^ and Notch^-ve^ cells is high **(Fig. 6B)**. Surprisingly, this region captured non-linear regimes, where tissue stretch begins only above a critical threshold of *pNotch* **(Fig. 6B)**. To understand whether our model recapitulates stretch dynamics in the compact layer, we fit it to an *in vivo* quantification of compact layer area at different levels of *pNotch*, measured using a *TP1:GFP* reporter **(Fig. 6C)**. Remarkably, we observed a distinct non-linear relationship between *pNotch* and compact layer area, with a weak correlation at low density, and a strong positive correlation at high density **(Fig. 6C’)**. The transition between these states occurred suddenly, at a threshold of ∼40% *pNotch* **(Fig. 6C’).** Our model therefore captures a non-linear response between *pNotch* and compact layer stretch. Instead of stretch occurring incrementally cell-by-cell, tissue stretch occurs rapidly after accumulation of a critical fraction of Notch^+ve^ cells. Thus, rapid morphological change is achieved through relatively small changes in the input signals around a critical threshold. This is reminiscent of phase transition like behaviour triggering spontaneous changes in tissue growth and mechanical properties observed in other developmental systems ^23,59–66^.

We next sought to understand the physical basis of non-linear tissue stretch. At a single cell level, our model displays a non-linear relationship between cell intrinsic tension and area, where cells stretch only when their tension falls beyond a tipping point, at which they yield to external forces **(Fig. 6D, S6D)**. Akin to a tug-of-war, the tipping point is where external force overcomes cell-intrinsic tension to allow for area expansion **(Fig. S6E)**. We then asked how this cell-intrinsic non-linearity manifests at the tissue-scale. Using a mean field simulation, we found that a non-linear tissue growth dynamic is lost when cells are uncoupled and vary freely in tension and area **(Fig. 6E**, *uncoupled*). However, a non-linear relationship between tissue growth and *pNotch* was restored by mechanically coupling cells, which we define as a convergence of forces experienced by different cells **(Fig. 6E**, *coupled*). In this coupled scenario, heterogeneity in cell tension and shape are averaged. And thus, cell stretch level depends on an effective tension, set by intrinsic properties and physical interfaces with surrounding cells.

When cells are mechanically coupled, our analysis suggests that Notch should act non-locally to lower effective tissue tension globally. In this scenario, cells would therefore collectively reach the tipping point required for them to yield to external force **(Fig. 6D)**, resulting in tissue-wide stretch. In support of this hypothesis, our vertex model shows that increasing *pNotch* increases the mean area of both Notch^+ve^ and Notch^-ve^ cells, in a non-linear manner **(Fig. 6F)**. Indeed, quantitative *in vivo* measurements showed that while stretch is biased towards Notch^+ve^ compact layer cells, all cells stretch in response to increasing *pNotch*, regardless of their Notch activation status **(Fig. S6F)**. Remarkably, using apicobasal thickness as a readout of stretch state, we further found that cells stretch more in neighbourhoods enriched with

Notch^+ve^ cells *in silico* and *in vivo*, regardless of their own Notch status **(Fig. 6G-H, S6G)**. These observations imply that mechanical coupling between cells allows mosaic Notch activation to globally decrease effective tissue tension. At a tipping point, tension falls sufficiently low that cells collectively yield to external force and stretch, resulting in rapid and uniform tissue expansion **(Fig. 6I)**. This critical transition thus efficiently converts variable cellular dynamics into robust organ-scale growth.

### Notch signalling triggers a morphological transition in myocardial surface area to tune ventricle size and pumping efficiency

Our theoretical modelling predicted that ventricle growth is triggered by a drop in the effective tension of compact layer cells to a tipping point, at which they collectively yield to external force. Fitting the model to *in vivo* measurements suggests this transition occurs between 72 and 96 hpf **(Fig. 6B, 7A)**. At 72 hpf, *pNotch* is too low to trigger tissue stretch, even if Notch^+ve^ cell tension is further depleted **(Fig. 7A**, dashed arrow). In contrast, the model predicts that ventricle size should be highly sensitive to changes in Notch^+ve^ cell tension at 96 and 120 hpf **(Fig. 7A’-A’’,** dashed arrows), when *pNotch* has crossed the critical threshold value of ∼40%.

**Figure 7.**
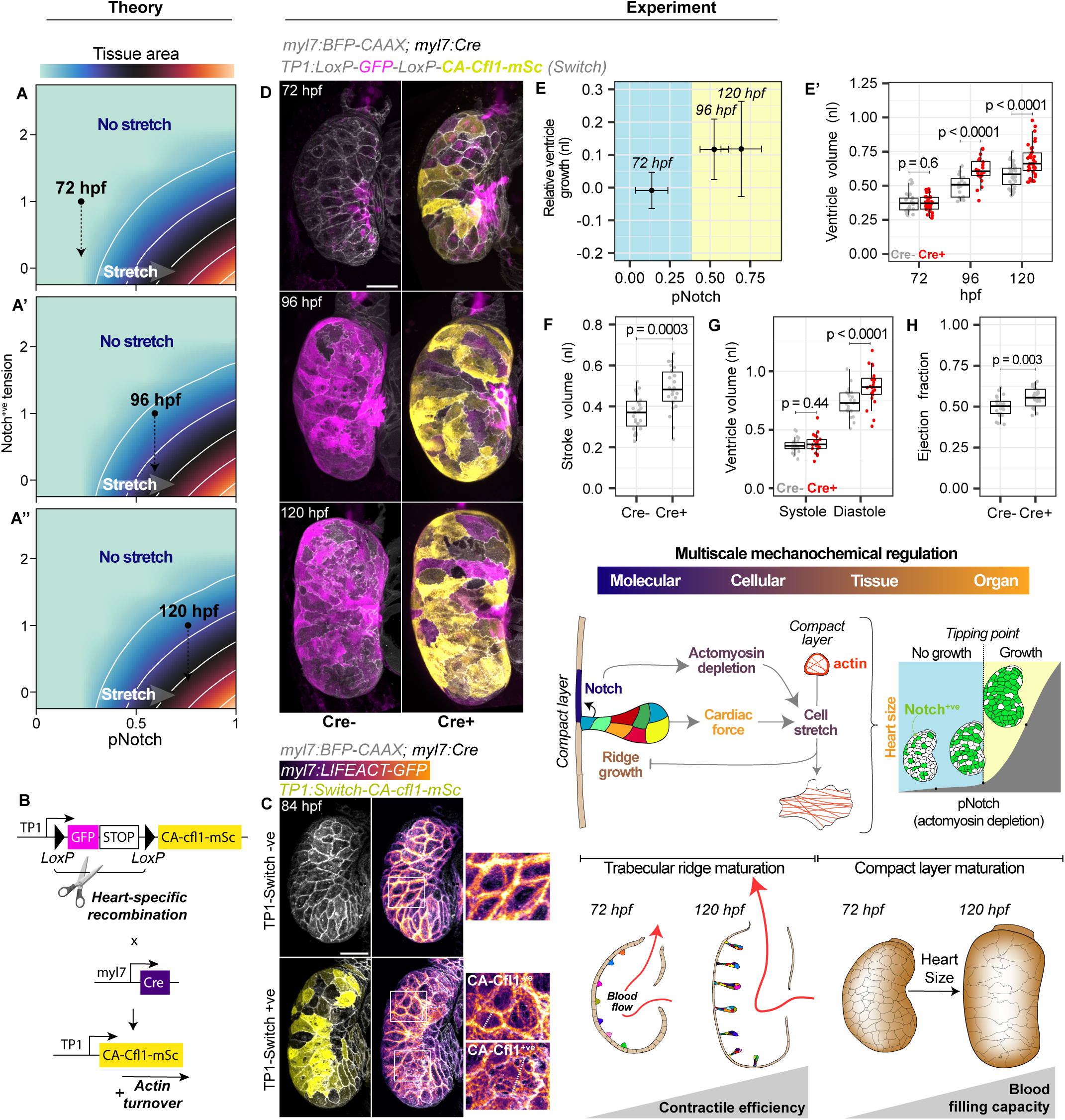
Notch signalling triggers a morphological transition in myocardial surface area to tune ventricle size and pumping efficiency. (A-A’’) in silico predictions for changes in tissue area in response to pNotch upon accelerated actin turnover in Notch+ve cells. Black dots show mapping of ventricles into model phase space at 72 hpf (A), 96 hpf (A’) and 120 hpf (A’’), with effect of experimentally reduced tension (dashed arrows). (B) Schematic for TP1:Switch strategy for targeting gene expression to Notch+ve compact layer cells. Cre excises GFP to drive Notch-dependent expression of CA-Cfl1 and accelerate actin turnover exclusively in compact layer cells. (C) CL surface projections showing actin (Lifeact) organisation in presence and absence of *TP1:Switch-CA-Cfl1-mScarlet*, with inlays (white boxes) showing actin in CA-Cfl1+ve and CA-Cfl1-ve cells. Representative of n = 14 embryos. (D) CL projections of *TP1:Switch-CA-Cfl1-mScarlet* ventricles in presence and absence of myocardial Cre, at 72, 96 and 120 hpf. (E, E’) Quantification of relative ventricle growth in *myl7:BFP-CAAX* (membrane); *TP1:Switch-CA-Cfl1-mScarlet* ventricles expressing myocardial Cre compared to Cre negative hearts, against proportion of Notch+ve cells measured using *TP1:GFP* reporter (pNotch), with error bars showing mean and standard deviation (E), or boxplots showing ventricle volume per condition per stage (E’). 72 hpf Cre-, n = 27; 72 hpf Cre+, n = 32; 96 hpf Cre-, n = 18; 96 hpf Cre+, n = 23; 120 hpf Cre-, n = 21; 120 hpf Cre+, n = 20. P values in (E’) show results of unpaired Students’ t-tests. (F-H) Quantification of stroke volume (F), end-systolic and end-diastolic volumes (G) and ejection fraction (H) for *TP1:Switch-CA-Cfl1-mScarlet* 120 hpf ventricles in the presence and absence of myocardial Cre. Cre-, n = 23 embryos; Cre+, n = 22 embryos. P values show results of unpaired Students’ t-tests. (I) Schematic summarising the multiscale mechanochemical feedback model driving morphological and functional maturation of ventricle. Scale bars = 30 µm

To experimentally test these model predictions *in vivo*, we devised a targeted genetic approach to disrupt actomyosin mechanics in Notch^+ve^ compact layer cells. We designed a Notch-sensitive *TP1:Switch* cassette, which drives expression of constitutively active Cofilin1 (CA-Cfl1)^67,68^ in Notch^+ve^ compact layer cells in the presence of myocardial Cre **(Fig. 7B-C, S7A-B’)**. CA-Cfl1 expression accelerated depletion of junctional actin only in compact layer cells **(Fig. 7C, S7B, B’)**, which we previously found to lower junctional tension and facilitate cell stretch **(Fig. 4A-D)**. In agreement with our model predictions, accelerating actin depletion in Notch^+ve^ cells with CA-Cfl1 failed to induce a significant change in ventricle volume at 72 hpf **(Fig. 7D, E, E’)**. However, at 96 and 120 hpf, where *pNotch* exceeds the critical threshold required for stretch (∼40%), expression of CA-Cfl1 invoked a significant increase in ventricle volume **(Fig. 7D, E, E’)**. Thus, ventricle size responds with qualitative similarity to our model, further supporting a non-linear growth regime where organ growth initiates rapidly through a critical drop in effective tissue tension.

Since heart function is dictated by its form^24,69,70^, we finally asked how regulation of ventricle size by cellular mechanics impacts cardiac function. With an experimental handle on ventricle size, we acquired timelapse movies of beating hearts expressing CA-Cfl1 and profiled changes in cardiac physiology **(Supplementary video 6)**. Remarkably, accelerated actin depletion in compact layer cells using CA-Cfl1 expression enabled a 31% increase in ventricle stroke volume **(Fig. 7F)**. This resulted from an 18% increase in ventricle filling capacity **(Fig. 7G)**, and an 11% improvement in ejection fraction **(Fig. 7H).** Thus, by accelerating junctional actin depletion in the compact layer cells thereby making them more deformable, we experimentally expanded ventricle size, and drove a supra-physiological improvement in its functional efficiency.

Together, our data support a multiscale model where molecular signalling, subcellular actomyosin remodelling, cell shape changes and organ function act in closed loop dynamics to drive robust morphological and functional maturation of the developing heart **(Fig. 7I)**. Trabecular ridges mature by recruiting cells from the outer compact layer. Trabecular maturation increases ventricle contractile efficiency and force production, which stretches compact layer cells and constrains excessive ridge growth. In parallel, stochastic Notch activity drives a global decrease in compact layer cell tension, triggering a rapid organ-scale growth required for optimal blood filling capacity.

## Discussion

A holistic understanding of how organ form and function emerge during development requires integration of feedback between microscopic processes, macroscopic deformations, and organ function^5,17,19,71^. Here, combining the excellent tractability of the zebrafish heart with quantitative approaches and controlled manipulations, we show that multiscale relay of mechanochemical information between the maturing compact and trabecular layers generates the functional tissue architecture of the vertebrate heart.

### Morphogenetic mechanisms building trabecular ridges

While genetic pathways underlying trabeculation are relatively well-studied^29,30,72,73^, cellular behaviours shaping trabecular ridges remain poorly understood. We now discover an autoinhibitory feedback loop that builds and constrains trabecular ridge density **(Fig. 7I)**. Trabecular ridges exhibit high degree of clonal heterogeneity in zebrafish as well as in mouse ventricles^27,74^. We now provide a cellular basis of this clonal heterogeneity in zebrafish and show that trabecular ridges expand by recruiting clonally distinct cells from the outer compact layer. By analysing relatively later stage of trabecular maturation at 10 dpf (days-post-fertilization)^36^, it was shown that TGFβ and Erbb2 signalling induced cardiomyocyte proliferation can promote trabecular growth. However, we find that total cell number increases by only 17% between 3-5 dpf zebrafish hearts. These results indicate that early-stage trabecular ridge growth is primarily driven by compact layer cell recruitment, while in the later stages, proliferation might be the dominant mechanism. Further, while migration or recruitment of compact layer cells have been hypothesised to contribute to trabecular growth in mice ^30,75^, our quantitative live imaging and a range of lineage tracing approaches provide detailed spatiotemporal insight into this process.

At the onset of trabeculation, as compact layer cells delaminate to seed the trabecular layer, adjacent cells activate Notch and are laterally inhibited from acquiring trabecular identity^25,26^. Using a stabilized Notch reporter^76^ to quantify persistent Notch activity in maturing trabecular ridges, we find that trabecular cells have never experienced Notch signalling **(Fig. S5E-E”)**. This suggests that ridges may mature by recruiting distant Notch-negative compact layer cells. Changes in the shape and position of trabecular seeder cells **(Fig. 1E)** may facilitate neighbour exchange to locate Notch-negative cells. Additionally, given the instructive roles of seeder or tip cells in other systems ^77^, seeder cells might actively organize ridges by exerting tensile forces on the compact layer cells, thereby recruiting them, akin to fly tracheal morphogenesis^78,79^.

Of note, we find that trabecular ridge growth is constrained by global cardiac forces, which stretch compact layer cells, thereby preventing them from entering the trabecular layer. Contrary to trabecular seeding ^25^, this control over ridge maturation by cardiac forces does not require Notch induced lateral inhibition. While increasing cardiac forces was sufficient to negatively regulate ridge density, this occurs independently of Notch activation **(Fig. S5D-D”)**. Because the level of cardiac force is governed by trabecular density **(Fig. 3A)**, trabecular morphogenesis should arrest in a self-limiting manner. The benefit of such feedback is clear given that the ridge density must be tightly controlled for optimal heart function^24,29,30,73^.

### Cell stretching as a strategy to trigger organ-scale growth and function

Because of the 3D curved geometry of myocardial wall, compact layer properties are not trivial to study and have been largely neglected until now^35^. Our bespoke morphometric analysis allowed us to examine unequivocally the properties of compact layer cells. We find that compact layer cells dilute their cortical actomyosin density and stretch, thereby enabling a rapid tissue expansion, despite losing cells to the trabecular growth. This extreme stretching of compact layer cells is reminiscent of superelasticity, a mechanism utilized by epithelial cells to undergo extreme deformations by diluting their cortical actin ^45^. However, unlike the reversible nature of superelasticity, compact layer cells undergo irreversible deformation over days, thus stably increasing myocardial tissue area. Cell stretching expands zebrafish retinal pigment epithelium^80^, enlarges the extra-embryonic region in avian embryos^81^, and massively increases the alveolar surface of the mouse lung^82^. While organ growth is typically linked to cell proliferation and hypertrophy, in these cases, cell stretch enables large-scale morphological transitions with limited proliferation or growth, which may reflect adaptation to nutritional and spatial constraints ^83,84^. Given the robustness of embryonic development, it is tempting to speculate that cell area increase has evolved as an efficient mechanism to induce fast tissue growth and establish organ architectures in the absence of sufficient time for cell proliferation.

As the heart develops, while the compact layer dampens its actomyosin cytoskeleton, the trabecular layer cells exhibit prominent sarcomere maturation. Although previous studies have indicated at these differences in vertebrate hearts^26,30^, we now provide a mechanistic and functional basis of these processes by selectively examining and perturbing compact layer mechanical properties. Our results suggest that the emergence of mechanically distinct adjacent tissue layers confers to the heart the duality of malleability and strength, necessary for its form and function. The compliant compact layer stretches to expand the heart size, which directly regulates how much blood the heart can pump. Meanwhile the inner contractile muscular trabecular ridges are the force generating component that propel the contractile efficiency of heart **(Fig. 7I)**.

### From cell-scale heterogeneity to organ-scale robustness

Developing tissues constantly buffer against stochastic fluctuations to preserve their architecture, but also rapidly change their macroscopic properties to grow and build complexity. These sudden morphological transitions, akin to phase transition dynamics, are non-linear in nature and are triggered by infinitesimal changes in molecular and cellular properties^50–52,85–89^. How these transitions are regulated and whether they have a functional role during organogenesis remains unclear^50–52,85^. Combining predictive theory, multiscale imaging and targeted perturbations, we show that myocardial tissue growth dynamics have putative signatures of a tissue phase transition. In this framework, cell-scale heterogeneity is converted into coherent organ-scale properties, resulting in rapid and stable myocardial growth that optimizes blood pumping efficiency. We find that the myocardial tissue transitions sharply from a non-stretched state to a stable growth regime once a threshold fraction of compact layer cells has activated Notch and diluted their junctional actomyosin. At this critical transition, global tissue tension drops sufficiently low that cells collectively yield to cardiac forces and stretch, resulting in organ-scale growth. Indeed, the spatial distribution of Notch activity is highly variable between embryos, but tissue-scale growth dynamics are highly conserved **(Fig. S6A-A’).** Thus, ventricle size is robust to inherently noisy cellular dynamics, but concerted changes at the cell-scale trigger stable transitions in organ size required to improve its function. These findings raise the intriguing possibility that critical transitions are general mechanisms regulating large-scale changes in cell behaviour during morphogenesis ^50,51,59,60,85–88^ and highlight the need to probe these transitions across scales to reveal the underlying regulation.

In conclusion, our study shows how robust functional morphology of a vertebrate heart emerges through multiscale mechanochemical coupling and form-function feedback. We anticipate that the regulatory mechanisms uncovered in this study will enable rational control of organ growth and potentially distil design rules building complex topological meshwork in other organs, for example lungs and kidney. Further, an integrated understanding of how developing embryos build complex 3D tissue structures and how those structures constrain organ function will deepen our understanding of developmental robustness, evolvability and identify novel mechanisms to engineer functional tissues *in vitro*.

## Supporting information

Supplementary modelling

## Author contributions

T.G.R.A. and R.P. conceived the project. T.G.R.A. designed and performed most of the experiments, generated the *TP1:Switch-CA-Cfl1-mScarlet* transgenic line, conceived the methods for image analysis and quantification, and analysed, visualised and curated the data. J.C-S. designed and analysed the 3D vertex model, with conceptual input from T.G.R.A. and R.P. M-C.R. generated the *myl7:Lifeact-mNG*, *myl7:H2B-mNG*, and *myl7:mScarlet-vcla* transgenic lines, performed all immunostaining and some chemical treatment experiments. K.G. generated the *TP1:mScarlet-PEST* transgenic line. J.B. supervised J.C-S. T.G.R.A. and R.P. interpreted the data and wrote the manuscript with input from all authors. R.P. supervised the project.

## Acknowledgements

We thank all members of the Priya Lab for helpful discussions throughout the project, and A. Yap, A. Elosegui Artola, A. Torres-Sánchez, C. Hill, D. Khoromskaia, G. Boezio, N. Tapon, S. Allanki, T. Frith and Z. Hadjivasoulou for critical reading of the manuscript. We thank S. Verger for guidance on using Surfcut2, and R. Fukuda for primer sequences to clone zebrafish Cofilin1. We thank R. Kelsh for gifting the *Zebrabow-M* transgenic line. We thank M. Renshaw and T. Fallesen from the Crick Advanced Light Microscopy facility for support with imaging and cell segmentation, and the Crick aquatics team for support with zebrafish husbandry. Work in R.P.’s laboratory is supported by the Francis Crick Institute, which receives its core funding from the Cancer Research UK (FC011160), the UK Medical Research Council (FC011160), the Wellcome Trust (FC011160), and the British Heart Foundation (SP/F/20/ 150014).

## Declaration of interests

The authors declare no competing interests.

## Supplementary Video Legends

**Video 1. Transitions in ventricle functional efficiency in wild type and trabeculation-inhibited embryos.**

Midsagittal timelapse movies of beating hearts at 72 hpf and treated with DMSO or PD168393 (PD) from 72 hpf to 96 hpf or 120 hpf. Videos acquired at 100 frames per second, playback at 20 fps.

Scale bar = 30 µm

**Video 2. Immediate effects of chemical perturbations on cardiac function.**

Midsagittal timelapse movies of beating hearts showing immediate effects of DMSO, 10 µM Nifedipine (NF), 100 µM Propranolol (PRO) and 50 µM IBMX, after 2-hours incubation from 80 hpf. Videos acquired at 100 frames per second, playback at 20 fps.

Scale bar = 30 µm

**Video 3. Immediate effects of chemical perturbations on cardiac function.**

Midsagittal timelapse movies of beating hearts showing immediate effects of distilled water or 50 µM Isoprenaline (ISO) after 2-hours incubation from 80 hpf. Videos acquired at 100 frames per second, playback at 20 fps.

Scale bar = 30 µm

**Video 4. Calcium transients in a beating heart.**

Midsagittal timelapse movie of 120 hpf beating heart showing dynamic fluoresence of *myl7:GCaMP*. Videos acquired at 100 frames per second, playback at 30 fps.

Scale bar = 30 µm

**Video 5. 4D beating hearts showing elastic deformation.**

3D timelapse movies of beating hearts at 72 hpf and 120 hpf, in 3D rendered view. Videos acquired at 100 frames per second, playback at 30 fps.

Scale bar = 30 µm

**Video 6. Functional effects of ectopic actin depletion on compact layer cells.**

Midsagittal timelapse movies of 120 hpf beating hearts expressing *TP1:Switch-CA-cfl1-mScarlet* in the presence and absence of *myl7-Cre*. Videos acquired at 100 frames per second, playback at 30 fps.

Scale bar = 30 µm

## Materials and Methods

### Zebrafish handling

*Danio rerio* were maintained at 28.0°C in fresh water (pH 7.5 and conductivity 500 µS) according to standard practice on a 15 hour on/9 hour off light cycle. All regulated procedures were performed under UK project license PP8356093 according to institutional (Francis Crick Institute) and national (UK home office) requirements as per the Animals (Scientific Procedures) Act 1986.

### Zebrafish transgenic lines and mutants

The following transgenic lines and mutants were used in this study: *Tg(myl7:BFP-CAAX)^bns193^*, abbreviated as *myl7:BFP-CAAX*; *Tg(myl7:EGFP-Hsa.HRAS)^s883^*, abbreviated as *myl7:HRAS-GFP*; *Tg(-0.8myl7:H2BmScarlet)^bns534^*, abbreviated as *myl7:H2B-mScarlet*; *Tg(-0.8myl7:H2B-mNeongreen)^fci600^* (this study), abbreviated as *myl7:H2B-mNG*; *Tg(myl7:H2B-EGFP)^zf521^*, abbreviated as *myl7:H2B-GFP*; *Tg(gata1a:DsRed)^sd2^*,abbreviated as *gata1:dsRed*; *Tg(myl7:Lifeact-GFP)^s974^*, abbreviated as *myl7:Lifeact-GFP*; *Tg(myl7:Lifeact-tdTomato)^bns141^*, abbreviated as *myl7:Lifeact-tdTomato*; *Tg(-0.2myl7:Lifeact-mNeongreen)^fci611^* (this study), abbreviated as *myl7:Lifeact-mNG*; *Tg(myl7:GgMylk-cpEGFP-RnCalm2)^s878^*, abbreviated as *myl7:GCaMP*; *Tg(-0.2myl7:mScarlet-vinculina)^fci619^* (this study), abbreviated as *myl7:mScarlet-vcla*; *Tg(myl7:actn3b-EGFP)^sd10^*, abbreviated as *myl7:actn3b-GFP*; *Tg(ubi:LOX2272-LOXP-Tomato-LOX2272-Cerulean-LOXP-YFP)^a131^*, abbreviated as *Zebrabow-M; Tg(cryaa:DsRed, -5.1myl7:CreERT2)^pd13^*, abbreviated as *myl7:CreERT2; nrg2a^mn0237Gt^*; *Tg(EPV.TP1-Mmu.Hbb:EGFP)^um14^*, abbreviated as *TP1:GFP*; *Tg(EPV.TP1-Mmu.Hbb:mScarlet-Mmu.Odc1)^fci613^* (this study), abbreviated as *TP1:mScarlet-PEST*; *Tg(Tp1-Hbb:hist2h2l-mCherry)^s939^*, abbreviated as *TP1:H2B-mCherry*; and *Tg(EPV.TP1-Mmu.Hbb:LoxP-GFP-LoxP-CA-Cfl1_S3A-mScarlet)^fci608^* (this study), abbreviated as *TP1:Switch-CA-Cfl1-mScarlet*. Cellular structures marked with fluorescent reporters are indicated in Table S1.

### Plasmid cloning

All plasmids generated in this study for transgenesis used a vector backbone (pTol2) that contains the Tol2 sites necessary for Tol2-mediated transgenesis ^90^. Briefly, unique pairs of restriction enzymes were used to linearize the pTol2 vector. Gene fragments to be cloned were either amplified by PCR using Phusion DNA polymerase (NEB M0530) and specific primers (Table S2) or synthesized (Twist Biosciences or Thermofisher). In-Fusion^®^ technology (Takara Bio #638947) was used to assemble the desired plasmids according to manufacturer’s protocol. All plasmids were sequence-verified using Sanger sequencing and/or full plasmid sequencing. The Lifeact-mNeonGreen synthesized fragment contained a zebrafish codon-optimised Lifeact sequence. All cloning strategies and plasmid maps are available upon request. The sequences of primers used for generating constructs used in this study are detailed in Table S2.

Wild type Cofilin1 was amplified with Taq RNA polymerase from a 48 hpf cDNA library, gifted by Luca Guglielmi (Caroline Hill Lab), using RT-PCR and the following primers: forward 5’ – ATGGCCTCCGGAGTTACAGTG -3’, reverse 5’ – GCCTCAATCGGTTAGAGGCTT – 3’. This fragment was then ligated into a pGEM-t EASY cloning vector. In-fusion primers including 5’ base substitutions were used to generate constitutively active Cofilin1. These primers were: CA-Cfl1 forward (5’ – GGTTCCGGATCGATAAGCTTGATATCACGCGTCCACCATGGCCGCCGGAGTTACAG – 3’), and CA-Cfl1 reverse (5’ – CCCTTGCTCACGGGTACCACCGGTCGATCGGTTAGAGGCTTTCCCTCC – 3’).

These sequences were based on a cloning strategy published previously^68^.

### Generation of genetic mosaics

Tol2 transgenesis^90^ was performed as described previously^25^. Briefly, zebrafish embryos were injected at the 1-cell stage with 1 nl of an injection mix containing 20 pg/nl Tol2 transposase, 5-20 pg/nl plasmid DNA and 1:10 phenol red. The amount of DNA injected for each construct was optimised using serial dilution. After injection, embryos were kept in 0.5X E2 (egg water) supplemented with 0.003% phenylthiourea (PTU, Sigma-Aldrich P7629) from 24 hpf and raised between 72 and 120 hpf for imaging. Injected embryos were screened for mosaic fluorescence at 72–96 hpf and imaged using confocal microscopy.

### Zebrafish transgenesis

Tol2 transgenesis was used to generate 5 new transgenic lines used in this study: *Tg(-0.2myl7:Lifeact-mNeongreen)^fci611^*, *Tg(-0.8myl7:H2B-mNeongreen)^fci600^*, *Tg(-0.2myl7:mScarlet-vinculina)^fci619^*, *Tg(EPV.TP1-Mmu.Hbb:mScarlet-Mmu.Odc1)^fci613^* and *Tg(TP1:LoxP-GFP-LoxP-CA-Cfl1_S3A-mScarlet)^fci608^*. Wild-type (WT) AB embryos were injected at the 1-cell stage with 1 nl of injection mix containing 20 pg/nl Tol2 transposase mRNA, 10-20 pg/nl plasmid DNA and 1:10 phenol red. F0 embryos were screened at 72 – 96 hpf, and those positive for fluorescence were raised to adulthood. Adult F_0_ (founder) fish were then outcrossed to WT animals, and their progeny (F_1_ generation) were screened for health and bright homogenous fluorescence either in the heart (*myl7*) and/or in known domains of Notch activity (*TP1*) and raised to adulthood. Further validations were performed by imaging F_2_ embryos in combination with other myocardium-specific transgenic lines. For each line, 3-9 founders were identified and only healthy transgenic line with single transgene insertion and homogeneous fluorescence was kept.

### Microscopy

Embryos were screened for positive fluorescence using a Leica SMZ18 fluorescence stereomicroscope from 72 hpf. They were then embedded in 1% (w/v) low melting point agarose with 2 µg/µl tricaine to immobilise the embryo and stop the heart, in a 35 mm diameter glass-bottomed dish. Once the agarose was set, the dish was filled with egg water containing 4 µg/µl tricaine. As stopping the heart causes it to collapse, all imaging was performed within 10 minutes of mounting unless stated otherwise. Most stopped-heart imaging was performed using a Zeiss LSM 710 or 980 Axio Examiner confocal microscope with a Zeiss W Plan-Apochromat 40x/1.0 DIC M27 water immersion dipping objective. A 1024 x 1024-pixel scan field was used at 1x optical zoom, yielding a pixel size of 0.21 µm. For faster imaging of large datasets acquired for wild type morphometric analysis, a Nikon W1 spinning disk confocal microscope equipped with a Nikon Apo LWD 40x WI λS DIC N2 water immersion dipping objective was used.

### Longitudinal imaging

For repeated imaging of the same embryo, imaging was performed as described above, but fresh egg water was added to the dish during imaging instead of egg water + 4 µg/µl tricaine. All imaging was performed within 10 minutes of stopping the heart. The embryo was then released from the agarose using forceps, and isolated in a 6-well plate in egg water. This process was repeated for each round of imaging.

### Photoconversion

Embryos with mosaic expression of *myl7:KikumeGR-NLS* were screened for bright mosaic fluorescence and mounted for longitudinal imaging at 72 hpf as described above. A nucleus from a single delaminating cell or trabecular seed was selected and a circular ROI was marked around it using the ‘Regions’ function in Zen Black. Photoconversion was performed exclusively in this ROI, using continuous acquisition, with a 405 nm laser set to 1-2% power (the same laser power used for routine membrane channel imaging). Photoconversion was performed for 1-3 seconds, until red fluorescence was strongly detected, and green fluorescence diminished. Embryos were then recovered and raised for subsequent imaging as described above.

### Timelapse imaging

For timelapse imaging, embryos were embedded in 1% (w/v) low melting point agarose, with 0.06 µg/µl tricaine to immobilise the embryo without disrupting the heartbeat, in a 35 mm glass bottomed dish. Embryos were then imaged using a Nikon W1 spinning disk confocal microscope equipped with a Nikon Apo LWD 40x WI λS DIC N2 water immersion dipping objective. All timelapse imaging was performed at a frame rate of 100 fps, with an exposure time of 10 ms. 2D slice movies were obtained using continuous acquisition for 10 seconds on a single midsagittal Z slice. For 3D timelapse imaging, 10-second movies were acquired sequentially for each Z-slice over a 70 µm Z-depth, with 1 µm slice spacing. Post-acquisition temporal registration was then used to align imaging data from each slice, to assemble a 4D movie^91^.

### Calcium imaging

*myl7:GCaMP* embryos were mounted for timelapse imaging as described previously, and 5-10 second movies were acquired on a single midsagittal section. A kymograph was generated using a spline oriented perpendicular to the myocardium using the ‘dynamic reslice’ function in FIJI, extending across the compact and trabecular layers and into empty space outside the ventricle and inside the ventricle lumen. Spline thickness was increased to 30 pixels to average signal from a broad region of myocardium. In the kymograph, separate splines were drawn through signal corresponding to the compact and trabecular layers, and used for a line quantification in FIJI.

### Pharmacological treatments

To inhibit trabecular growth, embryos were first manually dechorionated and then incubated in 10 µM PD168393 (Calbiochem #513033), or an equivalent volume of DMSO as vehicle control, in 50 ml petri dishes containing 25 ml PTU-treated egg water supplemented with 1% DMSO to aid drug solubilisation, at a maximum density of 50 fish. The modified media was replenished every 24 hours. For disruption of heart function, embryos were treated with 50 µM Isobutylmethylxanthane (IBMX; Sigma-Aldrich #15879), 50 µM Isoprenaline (Sigma-Aldrich #I5627), 10 µM Nifedipine (Sigma-Aldrich #N7634) or 100 µM Propranolol (Sigma-Aldrich #P0884). Stock solutions were prepared in DMSO for IBMX, Nifedipine and Propranolol, and distilled water for Isoprenaline. 8-12 embryos were treated with each drug in a total volume of 5 ml PTU-treated egg water supplemented with 1% DMSO in a 6-well plate from 80 hpf to 100 hpf. To inhibit Notch activity, embryos were treated with 2.5 µM LY411575 (Selleckchem #S2714), or an equivalent volume of DMSO as vehicle control. 8-12 embryos were treated with LY411575 in a total volume of 5 ml PTU-treated egg water supplemented with 1% DMSO in a 6-well plate. For actin stabilisation, embryos were treated with 500 µM Jasplakinolide (Invitrogen #J7473), or an equivalent volume of DMSO as vehicle control, in PTU-treated egg water supplemented with 1% DMSO.

### Whole-mount immunostaining

Immunostaining of zebrafish hearts was performed as described previously^25^. Deyolked embryos were treated with proteinase K (PK, Sigma #P6556) for 55 mins (72 hpf; 3 µg/mL), 40 mins (96 hpf; 6 µg/mL), or 50 mins (120 hpf; 6 µg/mL). The fin folds of 96 hpf embryos were cut using a scalpel and all three stages (72, 96, and 120 hpf) were combined in the same tube for immunostaining after PK permeabilization. 10% goat serum was used for blocking. Phosphorylated myosin light chain 2 primary antibody (Cell Signalling #3671) was applied at 1:125 dilution in combination with the goat anti-rabbit Alexa Fluor 555 secondary antibody (Life Technologies A21428; 1:500). To amplify myocardial membrane signal in fixed *myl7:BFP-CAAX* embryos, a BFP nanobody conjugated to Alexa 647 (2B scientific N0502-AF647-L; 1:125) was combined with the primary antibodies. Alexa Fluor 488 Phalloidin (Invitrogen A12379; 1:400) was applied during the secondary antibody incubation.

### Image processing

Generic image processing was performed in FIJI. Z-stacks were rotated where anteroposterior alignment deviated from the XY axis of the image using bicubic interpolation, and a median filter of radius 0.5 – 1 pixel was used for denoising where necessary. FIJI was also used for simple intensity quantifications using the analyze regions (local quantification of mean fluoresence intensity) and plot profile tools (quantification of intensity changes along a manually defined straight line or spline). No denoising was performed on signal to be quantified in downstream analysis. All 3D renderings were generated in Imaris (10.0.0) using the automated Surfaces pipeline.

### Nuclear detection and classification

Nuclear detection was performed in Imaris using either the Spots or Surfaces automated pipelines. In both cases, an expected object diameter of 3.5 µm was used for spot detection in the nuclear channel. A mean intensity filter for the nuclear fluorescence channel (ie. *myl7:H2B-mScarlet*, *myl7:H2B-mNeongreen*), was reduced until all nuclei were detected. Where the Surfaces wizard was used, a surface was generated across all nuclei, using a detail level of 0.5 µm, and then the Split Objects function was used to subdivide the surface between nuclear IDs. Spots detected in the atrium, valve and outflow tract were manually deleted to isolate ventricle nuclei. Nuclei were classified based on whether the corresponding cell was compact (in apical continuity with the outer myocardial layer), delaminating (in the compact layer, with elongated apicobasal length, reduced apical and expanded basal domains) or trabecular (discontinuous with the outer myocardial layer). Nuclei classified in each group were filtered into distinct object groups. 3D coordinates for nuclei in each group were exported in .csv format and pooled in an R script for quantification of cell number and spatial organisation.

### Ventricle superimposition and heatmap construction

Nuclear coordinates for individual ventricles were superimposed for population-scale spatial analysis using a modified Procrustes superimposition where each axis is scaled independently in length. Nuclei were first aligned to a standard coordinate system inferred for each ventricle. In brief, paired landmarks were placed in an orthogonal arrangement on a 3D surface of the ventricle in Imaris, marking a) the anterior and posterior, b) dorsal and ventral, and c) left and right limits. The 3D coordinates of each landmark were then exported with coordinates for all nuclei. In R, a new vector of points was inferred running between (and extrapolating beyond) each pair of landmarks, separated by 1 µm. The position of each nucleus along each axis was then calculated by selecting the point, from each axis independently, with the shortest Euclidean distance from the given nucleus. To aid superimposition, ventricle length along each axis was adjusted to the mean length measured across the whole population. Length was calculated as the maximum distance between nuclei along each axis. This range was normalised between 0 and 1, then multiplied by the mean for the whole population. In this method, each ventricle was allowed to scale freely on each axis, rather than enforcing isotropic scaling. Finally, coordinates from all nuclei, from all ventricles, were plotted in a common, normalised coordinate system. stat_density2d in the ggplot2 package was used to generate heatmaps for each cell state. A first layer was generated for compact layer nuclei that represents mean ventricle shape. A second layer was then added for delaminating or trabecular cells, showing variation in their spatial distribution in a ventral (XY) or anterior (XZ) view.

### Ventricle volume and surface area inference

Ventricle volume and surface area were calculated by fitting a concave hull to 3D nuclear coordinates for the ventricle. The Imaris Spots pipeline was used to measure 3D coordinate positions for all ventricle nuclei, excluding the outflow tract and atrioventricular canal, as described previously. Nuclear coordinates were then exported in .csv format and imported into R for downstream calculations. The ashape3d function in the alphashape3D package was used to infer a concave hull enclosing all coordinate points, which was visualised using the plot3d function from the rgl package. The alpha value in ashape3d was reduced until the surface tightly fit around all coordinate points, and then applied uniformly across all data acquired in the same experiment. The rvcg package vcgClean, vcgIsotropicRemesh and vcgSmooth functions were then used to clean and smooth the mesh. Volume and surface area measurements were acquired using the vcgVolume and vcgArea functions.

Where nuclear imaging was not possible (in 2D slice movies, and imaging without a nuclear reporter), ventricle volume was inferred by fitting a prolate ellipsoid to the dimensions of an object-oriented bounding box. Bounding box length was measured from the base of the outflow tract to the tip of the apex. Bounding box width was measured from the outer curvature to the atrioventricular canal. Volume, *v*, was then calculated as

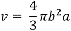

where *b* is the length of the short mediolateral semi-axis, and *a* is the length of the long anteroposterior semi-axis. This approach carries the assumption that ventricle is isometric on its mediolateral and dorsoventral axes.

### Clonal ID inference

Clonal IDs were inferred from imaging of the *Zebrabow-M* transgenic line. Female *Zebrabow-M* fish were outcrossed to male *myl7:BFP-CAAX; myl7:Cre-ERT2* fish. Embryos were treated with 10 µM 4-hydroxytamoxifen (4-OHT, Sigma-Aldrich H7904) from 60 to 72 hpf. Embryos were then washed 3 times with fresh egg water to dilute the 4-HT and prevent further recombination. Subsequently, embryos were incubated to 120 hpf and imaged with hearts stopped as described previously. For each embryo, fluorescence emission detected for each channel (CFP, YFP, RFP) was normalised to the maximum intensity detected in that channel. Quantification was then performed on single Z slices transecting each trabecular ridge. A circular ROI of 5 pixel diameter was used to locally sample normalised CFP, YFP and RFP fluorescence intensities for each cell in each ridge in each ventricle. For each ventricle, 5 control measurements were acquired in non-myocardial tissue. Clonal identity was then inferred based on the distance between cellular colour profiles in an RGB colour space. To calibrate this method, pairwise Euclidean distances were calculated between the 5 control ROIs for a given embryo, using the ‘dist’ function in R. The maximum distance between control points was defined as a uniqueness threshold, where any cells further apart than this threshold in colour space were considered unique in colour and thus clonally unique. Pairwise Euclidean distances were then calculated between each cell and all other cells in the same ridge. If the distance between a pair exceeded the uniqueness threshold, they were classified with the same clone ID. Calibration of the uniqueness threshold was performed independently for each embryo in the analysis.

### Surfcut

The Surfcut2^92,93^ plugin in FIJI was used for extraction of compact layer signal from Z-stacks of ventricles acquired using confocal microscopy. For pre-processing, Z-stacks were converted to 8-bit and resliced to a new slice thickness of 0.5 µm. The membrane channel (*myl7:BFP-CAAX* or *myl7:GFP-HRAS*) was input into Surfcut2 for binarization, edge detection and masking of the compact layer. Erosion depth was calculated independently for each dataset and developmental stage. A depth of 8-12 µm was used for 72-84 hpf ventricles, and reduced to 5-7 µm for 96-120 hpf ventricles where the compact layer is thinner. In each case, erosion depth was increased until signal from the trabecular layer started to appear in the maximum intensity projection, thereby ensuring good capture of the compact layer across its full apicobasal thickness. After calibration of Surfcut on a small sample of 3-4 embryos, mean parameters were applied for processing of all embryos in batch mode. All output Surfcut projections were manually examined to ensure no trabecular layer signal was included in the compact layer segmentation. Errors were corrected by repeating Surfcut processing for specific embryos in calibrate mode. Additional channels (ie. *myl7:Lifeact-mNG, myl7:mScarlet-vcla*) were processed using the masks generated for membrane signal in the same embryo. Output data were saved as maximum intensity projections for qualitative analysis, and Z-stacks for further downstream processing and quantitative analysis.

### Compact layer unwrapping

Compact layer signal extracted using Surfcut2 was unwrapped from the native 3D curvature of the ventricle onto a 2D plane for accurate measurements of cell geometry, using base functions in FIJI. Z-stacks of compact layer signal were opened in FIJI and rotated, using bicubic interpolation, for perfect alignment of the anteroposterior axis of the ventricle to the XY axis of the image. Where alignment required correcting in the ZY plane, the Z-stack was resliced from left, rotated, and then resliced from top to return to an XY view. In this orientation, Z-stacks were resliced from bottom for transversal sectioning through the ventricle. A maximum intensity projection was then generated for the central bulk of the ventral, excluding only the anterior and posterior poles. This central region of the ventricle was treated as a perfect cylinder for the purposes of tissue unwrapping. Using the maximum intensity projection, a spline was fit through the centre of the curved signal and spline thickness increased to cover all signal lying superficial or deep to the central curvature. This spline, representing the mean curvature of the ventricle across its anteroposterior extent, was saved to the ROI manager, and loaded onto the full transversal Z-stack. The straighten tool was then used to unwrap the curved signal into a 2D sheet, output again in transversal view. To return to an XY view, the straightened stack was resliced. Maximum intensity projections of unwrapped signal were then generated for all channels, and used for downstream 2D analysis of cell shape and cytoskeletal organisation.

### Manual myocardial segmentation

For segmentation of the trabecular layer, segmentation was performed using the manual surface construction workflow in Imaris. Imaging data was cropped to include only the ventral 50% of the ventricle. A spline with 0.5 µm point spacing was constructed every 2-3 slices (1 µm spacing) between the compact and trabecular layers on an xy sectioning plane, using fluorescent membrane signal. The resulting surface was used as a mask to isolate compact and trabecular layer signal for all channels.

### Cell segmentation

Unwrapped membrane projections were input into Cellpose ^94^ for 2D segmentation of compact layer cell apical domains. Predicted cell size and model match threshold were adjusted for best segmentation of the input data. Output label maps were then manually curated. Over-segmented cells, with multiple labels per cell, were merged. Under-segmented cells, with multiple cells within the same label, were deleted and manually redrawn. Cells drawn outside the ventricle, and in the atrium, valve or outflow tract were manually deleted. Cells were only retained if all boundaries with adjacent cells could be unequivocally resolved by eye.

### Quantification of cytoskeleton organisation

Segmentation label maps generated by Cellpose were merged with unwrapped projections of raw signal, including membrane and fluorescently-tagged or -labelled cytoskeletal proteins. The MorphoLibJ ^95^plugin in FIJI was used to quantify area values for each cell, and quantify mean signal intensity. For quantification of junctional and cytoplasmic intensity independently, labels were added to the ROI manager and scaled by a factor of 0.8 to generate cytoplasmic ROIs. Junctional ROIs were generated by subtracting cytoplasmic ROIs from the raw label map. Mean intensity values were then exported for each ROI set independently. Junctional: cytoplasmic ratio was calculated by dividing mean junctional intensity by mean cytoplasmic intensity. Multiple datasets were pooled after performing an independent min-max normalisation across all mean intensity values for each experiment.

### Signal orientation measurement

Local signal orientation was measured using the OrientationJ package in FIJI. The compact layer was isolated and flattened as described previously. A rectangular ROI was then isolated, representing the ventral surface of the ventricle. The quantitative orientation detection mode in OrientationJ ^96^was used to locally measure signal orientation in the *myl7:actn3b-GFP* channel, and the vector field mode was used to visually represent local orientations in the input image. Sarcomere orientation was inferred as lying perpendicular – with a 90° offset to – the actinin signal. For plotting, 0° was aligned to the horizontal, mediolateral axis, such that 90° aligned to the anteroposterior axis.

### Cardiac physiology measurements

Cardiac physiology measurements were acquired in FIJI based on dynamic changes in ventricle length and width measured in 2D slice movies of beating hearts. To calculate heart rate, a kymograph was generated using a spline through the ventricle apex. A straight horizontal line was then drawn that passed through the peaks in the kymograph. Fluorescence intensity was then measured along this horizontal line using the plot profile function, then the number of peaks was measured using the Find Peaks function in the *BAR* plugin. For 10 second slice movies, the number of peaks, reflecting individual heartbeats, was then multiplied by 6 to obtain heart rate in beats per minute (bpm). End-diastolic volume was inferred from ventricle length and width at 100% diastole, using the method described previously. Similarly, end-systolic volume was inferred at 100% systole. Stroke volume was calculated by subtracting end-systolic volume from end-diastolic volume. Ejection fraction was calculated by dividing stroke volume by end-diastolic volume. All volumes were calculated in µm^3^ and expressed in nl.

### Ventricle collapse assay

Embryos were mounted in 1% (w/v) low melting point agarose with 2 µg/µl tricaine for stopped-heart imaging as described previously. A timer was started as soon as the heart stopped, a first Z-stack was then acquired no more than 5 minutes later. Embryos were left mounted on the microscope until all successive imaging was completed, typically at 45 minutes after heart stop.

### Notch activity quantification

Fluoresence of *TP1:GFP, TP1:mScarlet-PEST* and *TP1:H2B-mCherry* Notch reporters was imaged as described above. To compare Notch activity with cell apical area, the compact layer was isolated and flattened as described above. In the flattened data, the fluoresence in the Notch reporter channel was binarized in FIJI. Cells were classified as Notch ‘on’ if their mode value was 255 in the binarized channel. To calculate Notch activity in 3D ventricles, nuclear segmentation was performed in Imaris, and nuclei were manually classified as compact or trabecular. Mean fluoresence intensity for the Notch reporter channel was then calculated in each nucleus. A blanket on/off threshold was calculated as the lowest mean intensity at which positive fluoresence could be observed relative to background. Where *TP1:GFP* was used to report Notch activity, segmentation of nuclear fluoresence was used to sample fluoresence in each cell. In all cases, Notch reporter activity was measured exclusively for the ventral half of the ventricle to abrogate error in on/off classification from signal attenuation on the Z axis.

### Vertex model summary

We established a 3D vertex model of the compact layer epithelium, considering apical (cortical and surface) cell mechanics, and taking into account cell volume preservation. The *in silico* tissue is modelled as a single layer of polyhedra, which are constrained to share an apical plane and have orthogonal lateral planes. To simulate tissue-wide stretch properties, we consider cells within a periodic square box, whose length acts as an additional degree of freedom in the model, and is our measure of tissue area. Cell mechanics are modelled by overdamped kinetics under the following energy functional:

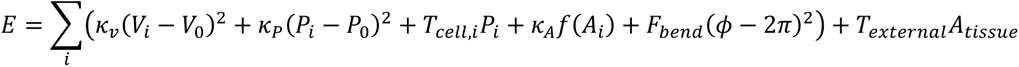

where each term corresponds to: (i) volume preservation; (ii) perimeter elasticity; (iii) cell cortical tension; (iv) surface elasticity/tension; (v) membrane bending minimization; and (vi) external force. We prescribe f(Ai) following ^56^ who propose apical surfaces have a constant surface tension at high areas, but below a critical apical area A0 show an elastic response, preventing apical areas from shrinking to zero. We prescribe each cell *i* with a Notch status (+/-) which specifies a distinct value of cell cortical tension *Tcell,i.* The energy is differentiated with respect to vertex coordinates and box size to obtain their respective velocities, and the system is solved to quasi steady-state, allowing T1 transitions to occur beneath a critical edge length as in^55 97^. The steady-state tissue area (L^2^ where L is the box size) and cell geometric properties are then calculated.

A grid parameter-scan was performed over the parameters: TNotch+, TNotch-, Texternal, pNotch. Linear regressions were performed between pNotch and tissue area, to obtain the extent of growth (slope) and the extent of non-linearity (sum of squared residuals). The scan was additionally fit to data on tissue area versus pNotch by evaluating the cost function:

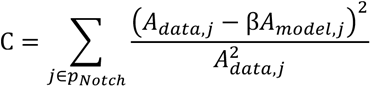

using a single free parameter β to rescale simulated tissue area to data units. The best-fit parameter combination is reported in Fig. 6B, C’ and used in Figs 6D, F, G’, S6D.

**Table S1.** Cellular structures/features labelled by transgenic fluorescent reporters.

| **Zebrafish line (common name)** | **Labelled structures/features** |
| --- | --- |
| *myl7:BFP-CAAX* | Cell membrane |
| *myl7:HRAS-GFP* |  |
| *myl7:H2B-mScarlet* | Cell nuclei |
| *myl7:H2B-mNG* |  |
| *myl7:H2B-GFP* |  |
| *gata1a:DsRed* | Red blood cells |
| *myl7:Lifeact-GFP* | F-Actin |
| *myl7:Lifeact-tdTomato* |  |
| *myl7:Lifeact-mNG* |  |
| *myl7:GCaMP* | Intracellular Ca<sup>2+</sup> |
| *myl7:mScarlet-vcla* | vinculin a |
| *myl7:actn3b-GFP* | actinin alpha 3b |
| *TP1:GFP* | Notch activity |
| *TP1:H2B-mCherry* |  |
| *TP1:mScarlet-PEST* |  |

**Table S2.**
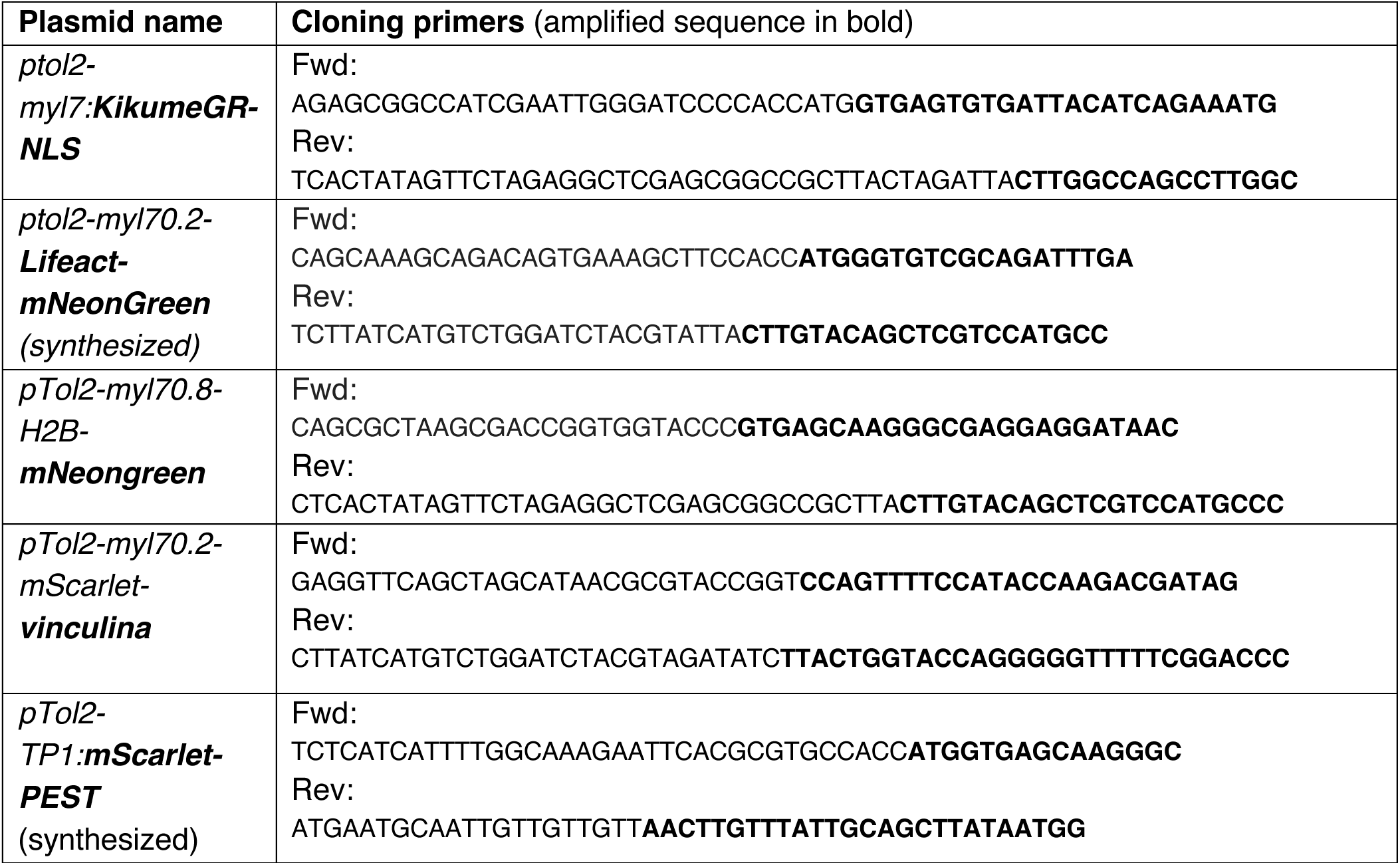
Primers used for gene cloning using In-Fusion^®^ technology.

## Notes

### Competing Interest Statement

The authors have declared no competing interest.

### Summary of Updates

Main figures moved down to prevent overlap with the stamper.

