## Supplementary figures for "Multiscale mechanics drive functional maturation of the vertebrate heart"

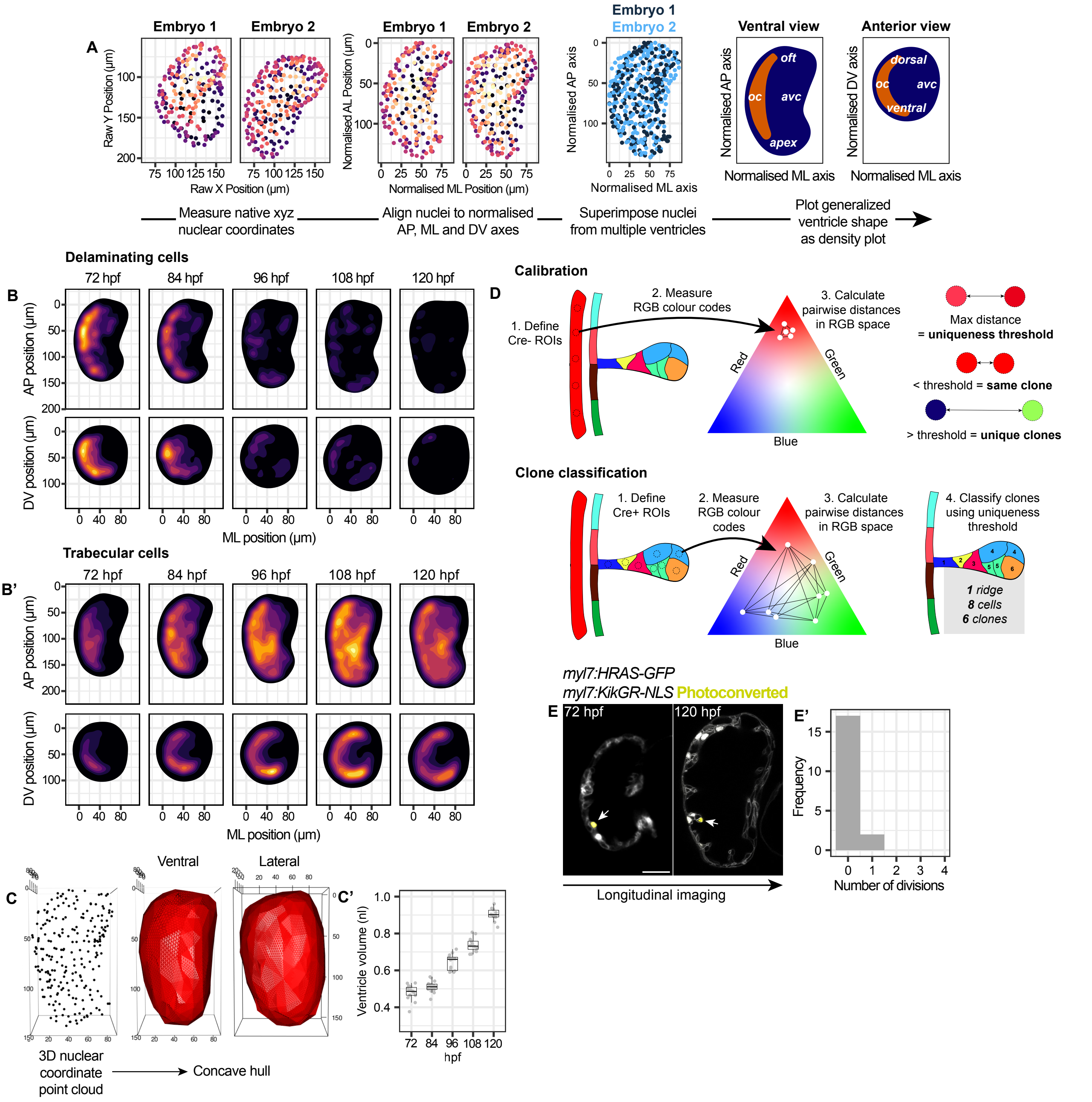


**Figure S1. Quantitative analysis of organ- and cell-scale dynamics of ventricle maturation, related to Figure 1**
(A) Pipeline for superimposition of cardiomyocyte nuclei for population-scale spatial analysis: raw nuclear positions, normalised nuclear positions, sample superimposition, density plots with spatial landmarks (outflow tract, oft; outer curvature, oc; atrioventricular canal, avc). (B, B') Density plots of (B) delaminating and (B') trabecular cell spatial positions generated for superimposed datasets for 72 - 120 hpf ventricles in ventral and anterior views using method in (A). n = 19,569 cells from 15 embryos per stage. (C-C') Example surface meshing from nuclear coordinates: raw coordinate point cloud for 120 hpf heart, surface mesh in ventral view and lateral view of outer curvature, and quantification of change in ventricle volume (C'). (D) Method for clone classification from Zebrabow-M imaging: calibration of clonal uniqueness threshold from Cre- ROIs, classification of clonal IDs from trabecular ridges using uniqueness threshold. (E-E') Midsagittal section of ventricle with mosaic expression of Kikume-GR-NLS, and trabecular seed photoconverted at 72 hpf, tracked longitudinally to 120 hpf (E) and histogram showing quantification of division events 72 – 120 hpf (E'). n = 17 tracked cells. Scale bars = 30 µm.

Figure S2. Andrews et al

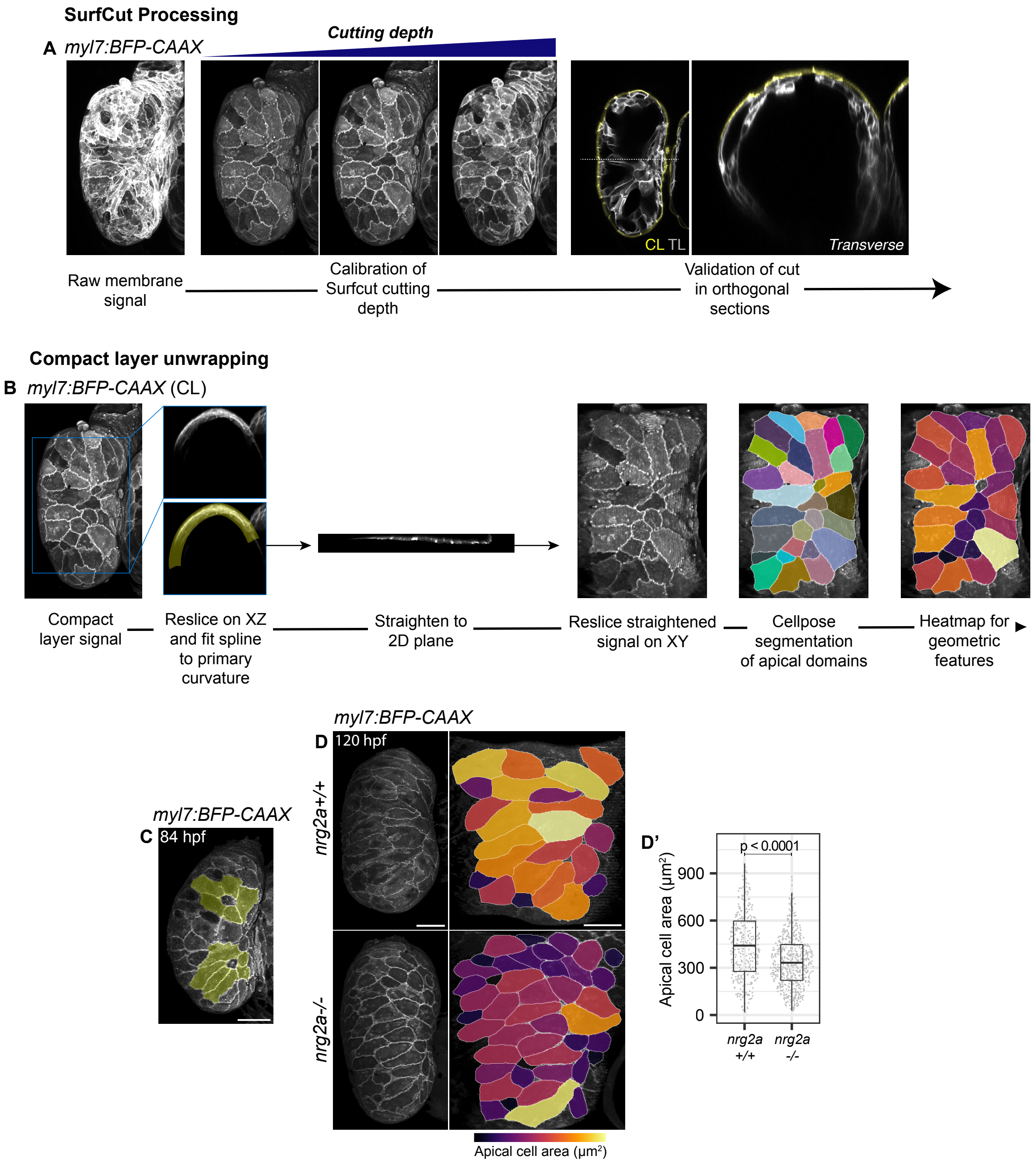


**Figure S2. Isolation, unwrapping and segmentation of compact layer cells, related to Figure 2.**
(A) Image processing pipeline for isolation of compact layer. Max projection of raw *myl7:BFP-CAAX signal*; surface projections of ventricle in (A) generated using SurfCut2 at increasing cutting depth, from shallow (apical signal only) to deep (trabecular signal visible in projection); validation of compact layer segmentation in orthogonal sagittal and transverse sections. (B) Image processing pipeline for unwrapping of compact layer signal. Transverse section of ventricle resectioned on XZ plane; spline fit to primary curvature of the signal; straightened signal; straightened signal resliced on XY to view the apical compact layer surface; Cellpose used for cell segmentation; geometric features are calculated using label maps; heatmaps are used to visualize cell-cell variations. (C) CL surface projections at 84 hpf with pseudocolour for cells in rosette organisation around delaminating cells. (D) CL projections and flat projections with cell area heatmap for ventricles of wild type and nrg2a homozygous mutant embryos, with quantification of compact layer cell area (D'). nrg2a+/+, n = 374 cells; nrg2a-/-, n = 624 cells. P value result of unpaired Mann-Whitney U test. Scale bars = 30 μm.

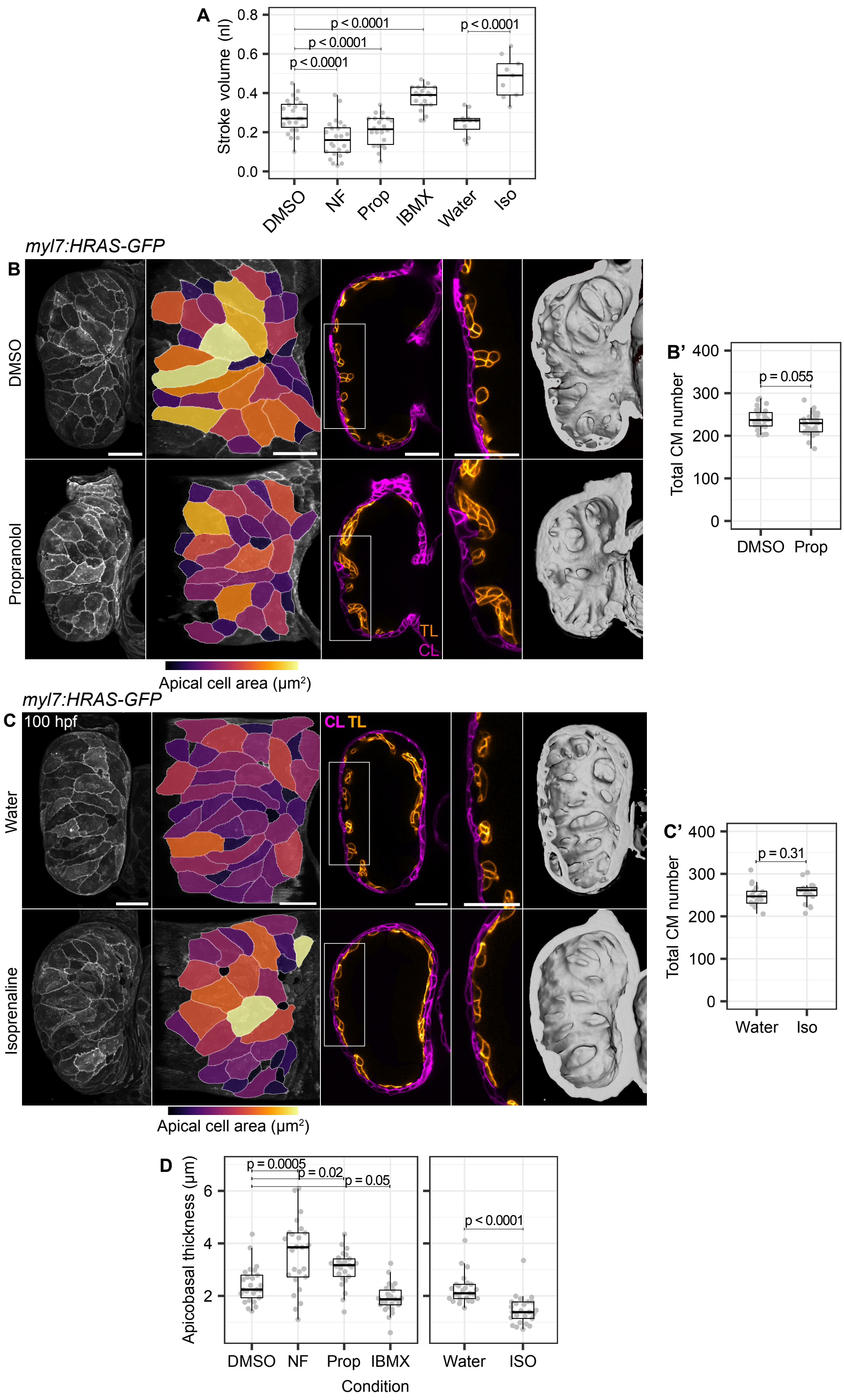


**Figure S3. Trabeculation drives compact layer cell stretch by increasing cardiac forces, related to Figure 3.**
(A) Measurement of stroke volume from slice movies of embryos treated with DMSO, 10 µM Nifedipine (NF), 100 µM Propranolol (Prop), 50 µM IBMX, water, or 50 µM Isoprenaline (Iso) for 2 hours from 80 hpf. DMSO, n = 24; NF, n = 24; Prop, n = 22; IBMX, n = 20; H2O, n = 12; Iso, n = 9. P values result of unpaired Students' t-tests. (B-C') Compact layer surface and flat projections with cell area heatmaps, midsagittal sections with pseudocolour for compact and trabecular layer, cropped panels for regions marked with white boxes, and 3D surface renderings of the trabecular layer, for ventricles of embryo treated with DMSO or Propranolol (Prop) (B), water or 50 µM Isoprenaline (Iso) (C) from 80 to 100

hpf, with quantification of total cardiomyocyte number (B’, C’). n = 30 embryos, DMSO; 34 embryos, Prop; 17 embryos, water; 20 embryos, Iso. P values result from unpaired Mann-Whitney U tests. (D) Quantification of compact layer apicobasal thickness in ventricles of embryos treated with DMSO, 10 μM NF, 100 μM Prop, 50 μM IBMX, water or 50 μM Iso from 80 to 100 hpf. P values show results of two-tailed unpaired Students’ t-tests. n = 25 measurements from 5 embryos per condition. Scale bars = 30 μm.

Figure S4. Andrews et al

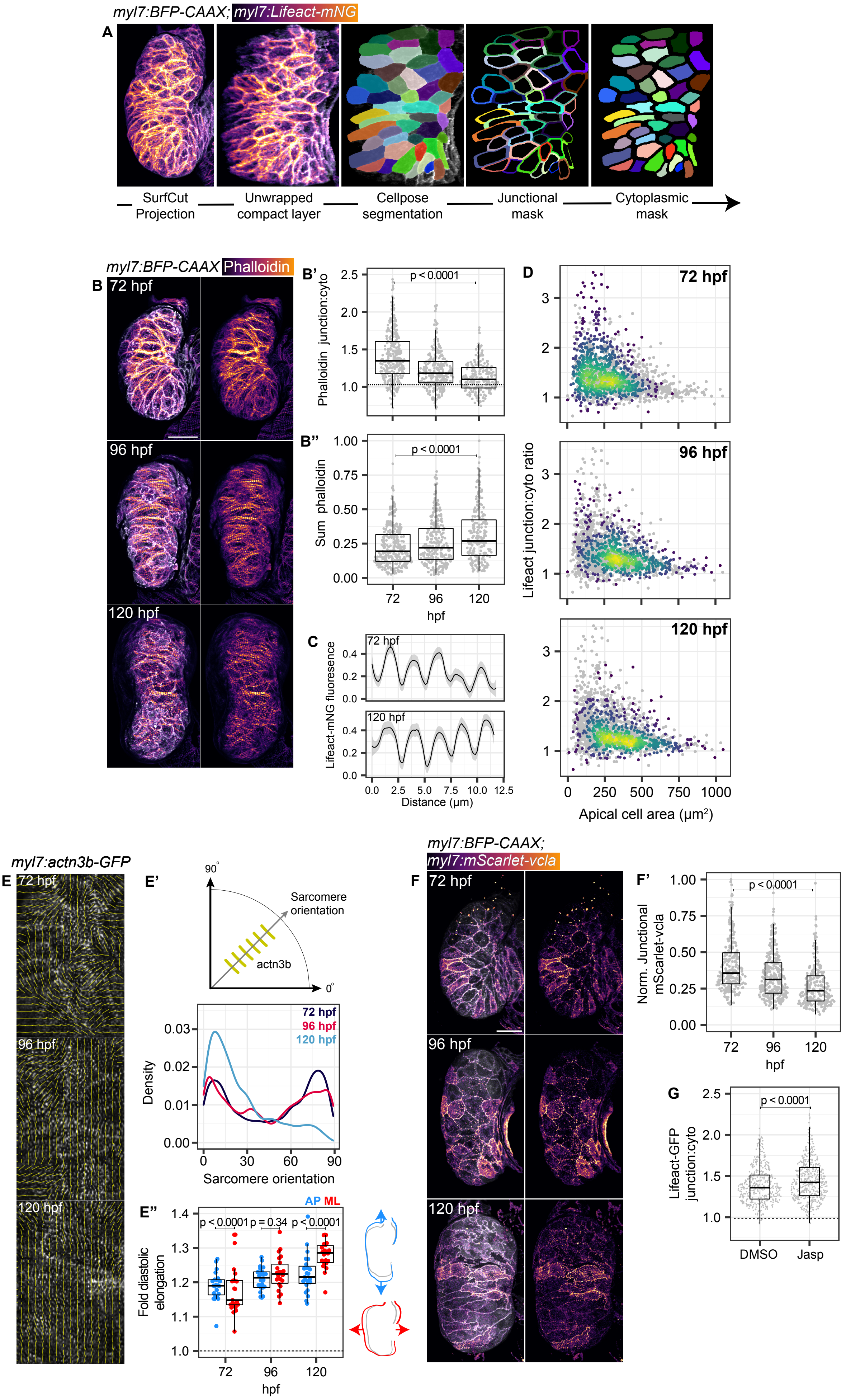

**Figure S4. Dynamics of actomyosin remodelling in stretched compact layer cells, related to Figure 4.**
(A) Image processing pipeline for measurement of junctional enrichment. CL surface projections unwrapped into a pseudo-2D sheet; cells segmented using Cellpose; junctional domains defined as the outer 20% of each label, cytoplasmic domains inner 80%. (B-B'') CL surface projections for fixed ventricles with phalloidin staining and detection of BFP-CAAX with FluoTag-X2 alexa fluor 647, with normalised quantifications of phalloidin junction:cyto ratio (B') and sum phalloidin signal per cell (B''). 72 hpf, n = 802 cells; 96 hpf = 498 cells; 120 hpf = 479 cells. P values show results of unpaired Mann-Whitney U tests. (C) Example line quantifications along actin cables visualized at 72 and 120 hpf with Lifeact-mNG fluoresence, indicating sarcomere length and frequency. (D) Correlation plots for compact layer cell apical area and actin (Lifeact-mNG) junction:cytoplasmic ratio, filtered for developmental stage. Heatmap shows point density. Grey background points show total dataset across all stages. 72 hpf, n = 757 cells; 96 hpf = 521 cells; 120 hpf = 541 cells. (E-E'') Local orientation of actn3b-GFP inferred with OrientationJ (E), with schematic and measurement of sarcomere orientation inference from actn3b-GFP orientation in a density plot (0 degrees mediolateral,ML; 90 degrees is anteroposterior, AP) (E'). 72 hpf, n = 11 embryos; 96 hpf, n = 10; 120 hpf, n = 13. Quantification of AP and ML diastolic stretch for embryos at 72, 96 and 120 hpf, with schematics (E''). 72 hpf, 20 embryos; 96 hpf; 21 embryos; 120 hpf, 21 embryos. (F, F') CL surface projections for ventricles showing intensity and organisation of mScarlet-vcla (F), with quantification of normalised junctional fluoresence (F'). n = 832 cells from 8 embryos, 72 hpf; 9 embryos, 96 hpf; 9 embryos, 120 hpf. P value results from unpaired Mann-Whitney U test. (G) Quantification of ventricle volume in embryos treated with DMSO or 0.5 μM Jasplakinolide (Jasp.) 80-100 hpf. n = 13 embryos, DMSO; 15 embryos, Jasp. P values result from unpaired Students' t-tests. Scale bars = 30 μm.

Figure S5. Andrews et al

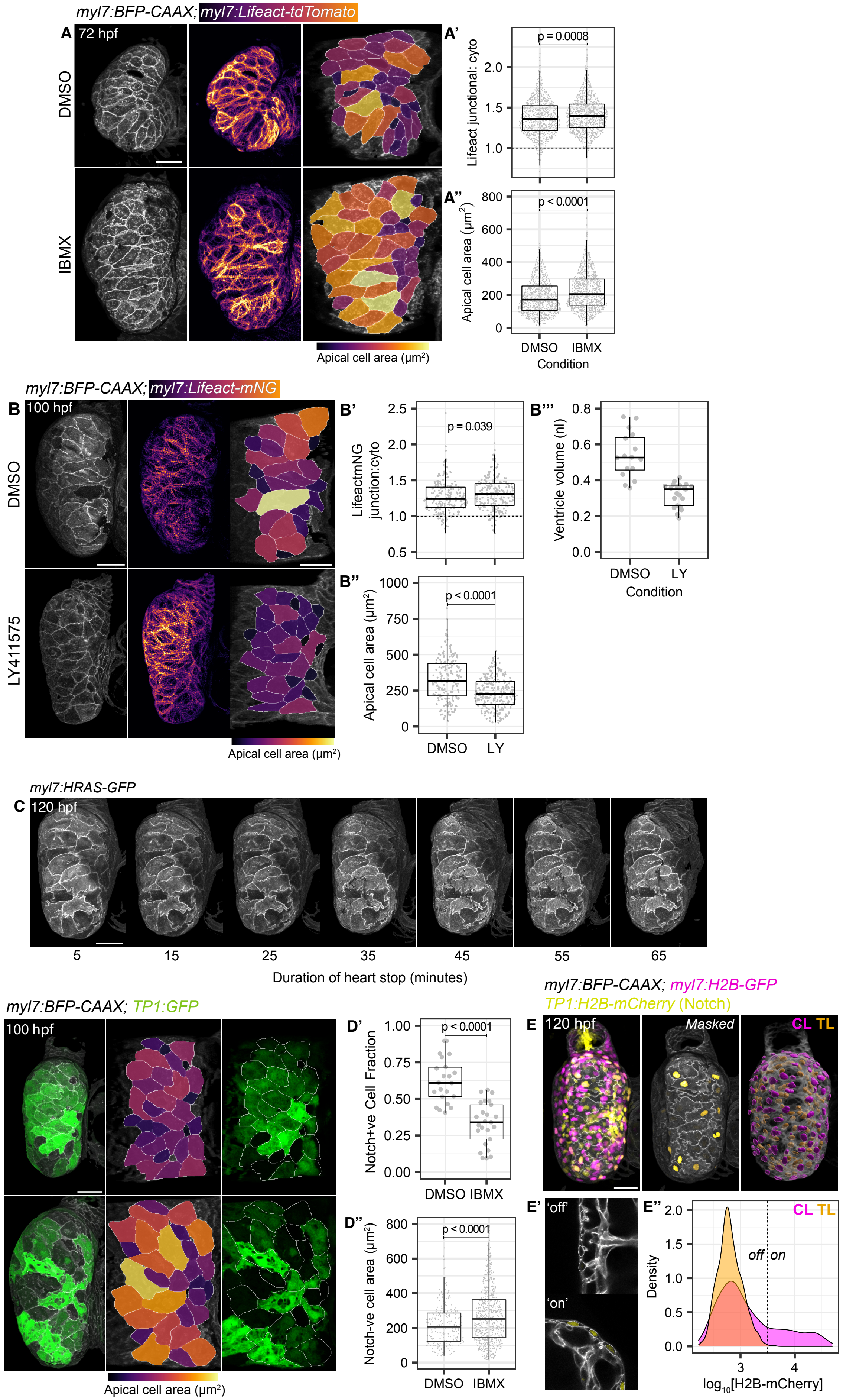

**Figure S5. Actomyosin remodelling facilitates compact layer cell stretch, related to Figure 5.**
(A-A'') CL surface projections and flat projections with compact layer cell area heatmaps for embryos treated with DMSO or 50 μM IBMX from 60 to 72 hpf (A), with quantification of junction:cytoplasmic ratio for Lifeact-tdTomato (A'), and cell apical area (A''). n = 913 cells, DMSO; 994, IBMX. P values result from unpaired Mann-Whitney U tests. (B-B''') CL surface projections and flat projections with cell area heatmaps for embryos treated with DMSO or 2.5 μM LY411575 (LY) 80-100 hpf (B), with quantification of Lifeact-mNeongreen junction:cytoplasmic ratio (B') apical cell area (B'') and ventricle volume (B'''). DMSO, n = 205 cells, n = 9 embryos; LY, n = 283 cells, n = 10 embryos. P values show results of unpaired Mann-Whitney U tests (B', B''), and unpaired Students' t-tests (B'''). (C) CL surface projections in longitudinal imaging of a 120 hpf ventricle collapsing following heart-stop in 4 μg/μl tricaine. (D-D'') CL surface projections and flat projections with cell area heatmap or cell outlines overlaid on *TP1:GFP* fluorescence, and quantification of Notch+ve cell fraction (D') and surface area of Notch-ve cells (D''). DMSO, n = 640 cells, 22 embryos; IBMX, n = 1052 cells, 25 embryos. (E-E'') CL surface projections of raw *myl7:H2B-GFP* and *TP1:H2B-mCherry, TP1:H2B-mCherry* masked using segmentation of the *myl7:H2B-GFP* signal (myocardial only) and 3D projection of segmented myocardial nuclei manually classified as compact or trabecular (E), sample *TP1:H2B-mCherry* fluoresence in myocardial nuclei, classified as 'off' or 'on' (E'), and density quantification for *TP1:H2B-mCherry* in compact and trabecular nuclei, relative to on/off threshold (dashed black line). n = 877 CL cells, 1177 TL cells, 9 embryos. Scale bars = 30 μm.

### Figure S6. Andrews et al

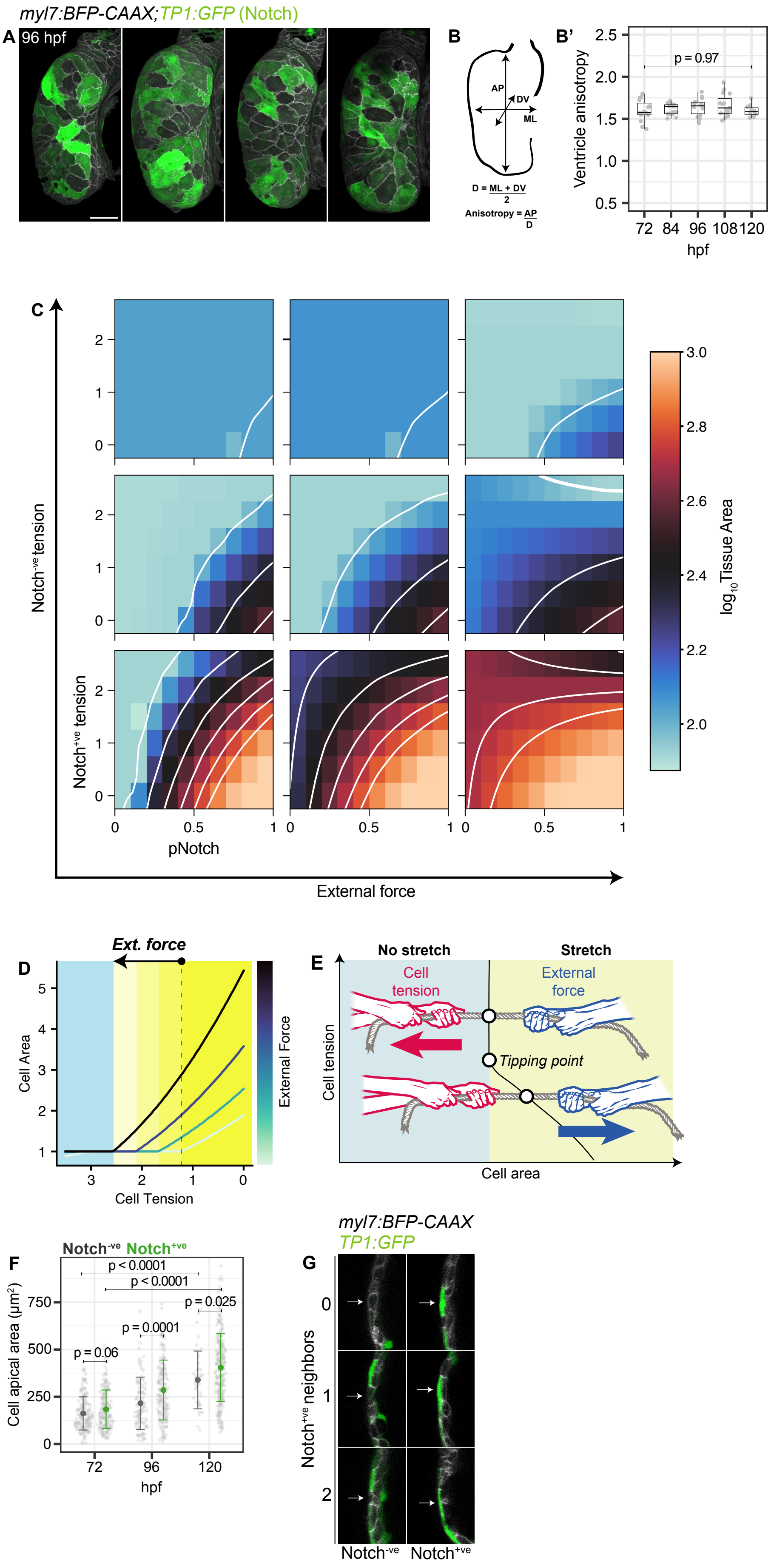

**Figure S6. Modelling of tissue stretch dynamics regulated by mosaic Notch activity, related to Figure 6**
(A) CL surface projections of four ventricles showing variation in *TP1:GFP* reporter fluoresence (Notch activity) between embryos. (B-B') Quantification of ventricle shape as relative length (anteroposterior, AP axis) and mean width (mediolateral, ML and dorsoventral, DV axes), termed ventricle anisotropy, schematic (B) and quantification between 72 and 120 hpf (B'). P value results from unpaired Students' t-test. n = 15 ventricles/stage. (C) 4D parameter scan for vertex model of tissue stretch. Sub-plots show heatmaps for tissue area with variation in pNotch and Notch+ve cell tension. Main plot shows variations in external force and Notch-ve cell tension. Solid white lines join regions with the same tissue area. (D) Effect of changes in external force on relationship between cell junctional tension and area. Changes in cell tension at the tipping point required for stretch are indicated by graded changes in blue (no stretch) and yellow (stretch) colour blocks. (E) Schematic showing force balance acting on single cells, inferred from the model. Cell-intrinsic tension acts to reduce cell area, while external force acts to increase cell area. When intrinsic tension is sufficiently high, cell area reduces to a non-stretched state. As tension drops, a tipping point is reached where tension is too low to resist cell area expansion driven by external force, resulting in cell stretch. (F) Quantification of compact layer cell apical area in Notch+ve and Notch-ve cells between 72 and 120 hpf, inferred using TP1:GFP reporter. 72 hpf, n = 371 cells, 8 embryos; 96 hpf, 298 cells, 8 embryos; 120 hpf, 300 cells, 10 embryos. Error bars show mean and standard deviation. (G) Cropped midsagittal sections showing thickness of example Notch-ve and Notch+ve cells in presence of 0, 1 or 2 Notch+ve neighbours inferred from *TP1:GFP* fluorescence. Representative of n = 85 Notch-ve cells from 18 embryos and n = 80 Notch+ve cells from 18 embryos.

Figure S7. Andrews et al

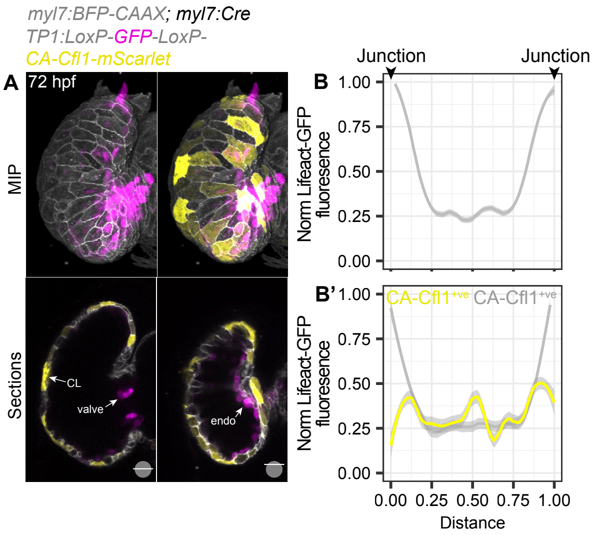


**Figure S7. TP1:Switch-CA-Cofilin1 strategy to dissociate actin in compact layer cells, related to Figure 7**
(A) Maximum intensity projections and sagittal sections showing tissue-specific expression of *TP1:Switch-Cfl1-mScarlet* at 72 hpf (CL, compact layer; endo, endocardium). Schematics in bottom right of each section indicate Z depth of the section. (B-B') Line quantifications across wild type (B), CA-Cfl1-ve and CA-Cfl1+ve cells (B') following dashed lines in (Fig. 7C inlays). Representative of n = 14 embryos.
