## Supplementary modelling for "Multiscale mechanics drive functional maturation of the vertebrate heart"

#### 1 3D Vertex Model Overview

To understand how mosaic activation of Notch leads to the growth of the zebrafish heart, we established a 3D vertex model of the compact layer epithelium [Farhadifar et al., 2007, Bielmeier et al., 2016, Misra et al., 2016, Alt et al., 2017]. In brief, this model considers apical (cortical and surface) mechanics of cells, as well as taking into account cell volume preservation. Further, the tissue is modelled using 2D (apical plane) periodic boundary conditions, with free boundary conditions in the z-plane. The periodic square box in which the tissue is situated (parameterised by the box length $L$) is itself a variable in the model, allowing for us to model tissue growth, without introducing artificial boundary conditions, capturing the idea that all cells of the epithelium are pulling/pushing on a common surface to drive global organ growth (while also taking into account local cellular rearrangements and deformations) [Guerrero et al., 2019, Bocanegra-Moreno et al., 2023].

##### 1.1 Cell geometry

Geometrically, cells are defined as polyhedra. Specifically, 3D cell volumes are constructed by considering 2D polygons – parameterised by a set of vertices – on the apical surface, and projecting these 2D points basally (i.e. a third degree of freedom for every vertex). This enforces that basal vertices of neighbouring cells meet a single points, and also assumes that the lateral surfaces of cells are perpendicular to the apical plane (which from our observations is a reasonable assumption). From this we can calculate, for each cell $i$, the apical perimeter $P_i$, the apical surface area $A_i$, the apical edge length of a pair of cells $\ell_{ij}$, and a cell's volume $V_i$. Further, given that in three dimensions, a polygon of more than three points need not lie on a single plane, we assume that there exists some bending energy on the basal surface to encourage it to be flat, parameterised by quantifying the residual curvature of the basal surface $\phi_i$. Details can be found in Appendix A.

##### 1.2 Energy and dynamics

Temporal evolution of the tissue is modelled by overdamped kinetics, underpinned by an energy functional that we prescribe to be:

$$E = \sum_i \left(\kappa_V(V_i - V_0)^2 + \kappa_P(P_i - P_0)^2 + T_{cell,i}P_i + \kappa_A f(A_i) + F_{bend}(\phi - 2\pi)^2\right) - T_{external}A_{tissue} \quad (1)$$

where

$$f(A) = \begin{cases} (A - A_0), & \text{if } A > A_0 \\ (A - A_0)^2 & \text{otherwise} \end{cases} \quad (2)$$

following the 3D vertex model from [Bielmeier et al., 2016], which proposes that apical surfaces have a constant surface tension at high areas, but below a critical apical area $A_0$ show an elastic response, preventing apical areas from shrinking to zero. In the energy functional, this is scaled by a pre-factor $\kappa_A$ in both cases to result in a continuous function in $E$ for varying $A$, with a minimum at $A = A_0$.

The parameters are:

- $\kappa_V$ - bulk modulus of volume deformations
- $V_0$ - preferred volume
- $\kappa_P$ - elastic modulus of perimeter/cortex
- $P_0$ - preferred perimeter (taking into account cortical tension and adhesive contributions)
- $A_0$ - preferred area
- $\kappa_A$ - surface tension / elasticity.
- $T_{external}$ - external force applied to the tissue (simulating fluid pressure)
- $F_{bend}$ - bending energy of the basal surface
- $T_{cell,i}$ - the junctional tension of the cell $i$

$\kappa_V$ and $F_{bend}$ are chosen to be large with respect to the other parameters to satisfy our observations that shape change is volume preserving, and that the basal surfaces of cells are flat. We use a minimalistic description of the external force with the term $T_{external}A_{tissue}$, such that the force is isotropically applied to the surface.

The 3D vertices comprising a cell can be defined by the 3D positions of basal vertices $\boldsymbol{x}_{\boldsymbol{j}}^{b}$ (and their corresponding projections onto the apical plane $\boldsymbol{x}_{\boldsymbol{j}}^{a}$; see Appendix A.1), as well as the size of the box. We choose a normalisation strategy for vertex coordinates so that local cell rearrangements versus global changes in tissue size (i.e. periodic box size) can be simulated in parallel. Let the coordinates of $\boldsymbol{x}_{\boldsymbol{j}}^{b} = (x_j^b, y_j^b, z_j^b)$. For convenience, let $L = \ell L_0$, where $L_0$ is the box size at the start of the simulation. Then we can define the relative 2D coordinates of vertices in terms of $x_j^b = \ell L_0 \hat{x_j}^b; y_j^b = \ell L_0 \hat{y_j}^b$. To aid numerical stability, given the 3D components of the model have larger parameter values (making this a 'stiff' problem), we choose a rescaling of $z_j^b$ such that changes in $\ell$ are iso-volumetric, effectively enforcing the inevitable fast-timescale relaxation of cell volumes to their optima upon global tissue stretching. Specifically, we choose $z_j^b = \frac{L_0}{\ell^2} \hat{z}_j^b$.

From this, we can recast the parameters of the model in terms of these effective, normalised variables, and calculate separately $\partial E/\partial \hat{x_j}^b, \partial E/\partial \hat{y_j}^b, \partial E/\partial \hat{z_j}^b, \partial E/\partial \ell$ to solve the dynamics of the system. This is equivalent to:

$$\frac{\partial \boldsymbol{x}_{\boldsymbol{j}}^{b}}{\partial t} = -\mu_x \frac{\partial E}{\partial \boldsymbol{x}_{\boldsymbol{j}}^{b}}. \tag{3}$$

and

$$\frac{\partial \ell}{\partial t} = -\mu_L \frac{\partial E}{\partial \ell} \tag{4}$$

##### 1.3 Simulation details

We initialise the tissue with a 2D voronoi diagram that approximates an equal-sized hexagonal lattice plus some noise in the initial conditions (81 cells). By inspecting $E(L)$ setting $F_{bend}$ and $\kappa_V$ to 0 (i.e. assuming that the associated features of cell geometry quickly reach equilibrium), we optimise $L$ to minimise the initial energy of the lattice, thus choosing a sensible scale to start the simulation. We then project basal vertices to equal distances such that the total volumes of cells are fixed at $V_0$. As these projections are equal and by definition perpendicular to the apical plane, $\phi = 0$.

Vertex dynamics are evolved in time using the forward Euler integration strategy, using JAXlib [Bradbury et al., 2018] to calculate the energy gradients. The simulation is solved until $t = 1000$, using a time-step of $\delta t = 0.02$.

We model the effect of Notch on cell mechanics as a reduction in the cell cortical tension $T_{cell,i}$. Specifically, cells of a particular Notch status are assigned the same parameter. Given a value of the frequency of Notch positive cells, $p_{Notch}$, the number of Notch positive cells is chosen as the nearest integer to $n_{cell}p_{Notch}$, and Notch status is allocated randomly to the cell ensemble. All other cell parameters are kept constant.

After simulation completion, we can calculate tissue area as $(\ell L_0)^2$, as well as cell geometric properties including area and basal extent.

##### 1.4 Parameter screen

We simulated our model over four parameter dimensions ($T_{cell}^{Notch+}, T_{cell}^{Notch-}, T_{external}, p_{Notch}$), across a $10 \times 10 \times 10 \times 10$ grid, and for each parameter combination, we simulate the model across 10 random seeds (affecting the positioning of Notch positive cells, and the minor random noise in the deviations of the initial vertex positions from a regular hexagonal tessellation). This gives a total set of $100,000$ simulations (Fig. S6C). Given symmetry in the parameterisation, we simulate only scenarios where $T_{cell}^{Notch+} \leq T_{cell}^{Notch-}$ and fill in the unsimulated parameter-combinations by a change of labels.

Our parameter choices were:

| | |
|---|---|
| $T_{cell}^{Notch+}$ | 0-0.027 (inclusive, 10 linear steps) |
| $T_{cell}^{Notch-}$ | 0-0.027 (inclusive, 10 linear steps) |
| $p_{Notch}$ | 0.05-0.95 (inclusive, 10 linear steps) |
| $T_{external}$ | 0-0.05 (inclusive, 10 linear steps) |
| $V_0$ | 1.0 |
| $\kappa_A$ | 0.02 |
| $\kappa_P$ | 0.003 |
| $\kappa_V$ | 0.3 |
| $A_0$ | 1. |
| $P_0$ | 4.5 |
| $F_{bend}$ | 0.1 |

Table 1: Table S1: 3D vertex model parameters, across 4D parameter screen

Note that the perimeter terms of the energy can be written as [Bi et al., 2015]:

$$E_{perim} = \sum_i \left[\kappa_P (P_i - P_0)^2 + T_{cell,i} P_i\right] \tag{5}$$

$$= \sum_i \kappa_P (P_i - \underbrace{(P_0 - \frac{T_{cell_i}}{2\kappa_P})}_{=P_{0,i}^{eff}})^2 - \underbrace{(P_0 - \frac{T_{cell_i}}{2\kappa_P})^2}_{=\text{constant}} \tag{6}$$

Therefore, in our code, we vary $P_{0,i}^{eff}$, with $T_{cell,i} = 0$. Above we report the effective value of $T_{cell,i}$ holding $P_0$ fixed; in practice, $P_0^{Notch+/-}$ is varied linearly in 10 intervals between 0 and 4.5.

#### 2 Mean field model

To understand what drives the non-linearity in tissue growth as a function of $p_{Notch}$ we applied some approximations to the model to make it analytically tractable.

##### 2.1 Mechanics of a single cell

We start by considering the mechanics of a single cell in the model. Let a cell's geometry be defined by a single parameter $A$, defining the (apical) area of a cell described by a regular hexagon. From this, we can calculate its edge-length $l = \sqrt{\frac{A}{3\sqrt{3}/2}}$ and its perimeter $P = 6\ell$. Considering the slower dynamics in the model (i.e. assuming volume preservation), and ignoring basal curvature, we can simplify the energy of a cell (embedded in a homogeneous tissue) as:

$$E = \kappa_P (P - P_0^{eff})^2 + \kappa_A f(A) + T_{external} A \tag{7}$$

where

$$P_0^{eff} = (P_0 - \frac{T_{cell}}{2\kappa_P}) \tag{8}$$

Given this equation, we can numerically solve for $\partial_A E = 0$ to define the steady state area $A^*$ as a function of the model's parameters. What this revealed is a non-linear relationship between $A^*$ and

$T_{cell}$: at high areas, cell area falls linearly with cell tension, whereas when cell area reaches a critical point, the area becomes much less compressible (Fig. S6E).

To test the effect of external forces (which in this reduced model implicate both the external forces applied on the tissue in the form of e.g. fluid pressure, but also the local forces exerted by other cells in the tissue), we varied $T_{external}$ as a proxy for these combined effects finding that the critical cell tension at which cells become quasi-incompressible shifts as a function of this external force (Fig. S6D).

#### 2.2 Examining the size of the tissue given mosaic Notch activation

We tried to connect this cell intrinsic property of cell stretch as a function of its internal and external forces, to the global process of tissue growth via a toy model that acts as a useful null hypothesis. Supposing every cell follows its own energy functional, uncoupled from one another, as defined above, and that Notch positive cells have a lower $T_{cell}$ than Notch negative cells, then cells of each Notch status would have their own optimal areas $A^+$ and $A^-$. From this, the area of a tissue with $N$ cells would be:

$$A_{tissue} = N(p_{Notch}A^+ + (1 - p_{Notch})A^-) \tag{9}$$

Which simplifies to:

$$A_{tissue} = N(A^+ - A^-)p_{Notch} + NA^- \tag{10}$$

This would imply that irrespective of the values of $A^{+/-}$, tissue area should scale linearly with $p_{Notch}$. Indeed this result holds true regardless of the energy functional or parameters chosen. This is at ends with our observations in the embryo or our simulations.

We posit that what is missing is the inherent coupling of mechanics between cells within an epithelium. This coupling comes in multiple forms: (a) local cell contacts, and the meeting of cell edges at common vertices, means that the stretching of one cell has an influence on another; (b) heterogeneous tensions due to mosaic Notch activation means that cells experience different local tension profiles depending on their positioning and local Notch density; (c) given cells are embedded in a common epithelium, a surface that has the capacity to stretch and deform, a given cell can experience the forces exerted by other cells at long-range; (d) if cell volume is preserved as a strong constraint, and basal surfaces show low curvature, there is an effective penalty on local apical area variance. Indications of this mechanical coupling emerge from analysis of the equations of the full model (Appendix B). We model these multifarious contributions in a simplified manner by stating that as cells become maximally coupled, the effective forces experienced by cells and therefore their sizes should converge.

Thus we consider an energy functional of a tissue that follows:

$$\begin{aligned} E = \kappa_A(p_{Notch}f(A^+) + (1 - p_{Notch})f(A^-)) + T_{external}(p_{Notch}A^+ + (1 - p_{Notch})A^-) \\ +\kappa_P \left(p_{Notch}(P^+ - P_0^{eff+})^2 + (1 - p_{Notch})(P^- - P_0^{eff-})^2\right) \\ +\alpha(A^+/A^- - 1)^2 \end{aligned} \tag{11}$$

where $\alpha$ defines the coupling strength. This model has two degrees of freedom $A^+$ and $A^-$, the areas of Notch+ and Notch- cells, where $P^+$ and $P^-$ are calculated as above. What this demonstrates is that as coupling increases, there is an increasing divergence from a linear response of tissue area to $p_{Notch}$. In the non-linear, coupled regime, the global response of tissue area to $p_{Notch}$ tracks the response of individual cells to cell tension; i.e. coupling allows for changes in $p_{Notch}$ to alter the effective tension experienced by cells, and therefore their resultant areas.

### Appendices

#### A 3D Vertex model

##### A.1 Geometric construction

In the main text, we present a 3D vertex model of the compact layer of the zebrafish heart. Cells are modelled as polyhedra, defined by a set of vertices. Specifically, we enforce cells to exist in an single-cell layered epithelial-like sheet, where further, the apical surfaces of cells are coplanar. If we additionally assume that the lateral surfaces of cells are orthoganal to the apical plane, we can describe each cell by a set of vertex positions in 3D.

Let $i$ denote the cell index and $q$ denote the vertex index. Given orthoganality between apical and lateral surfaces, the basal vertices of each cell ($\mathcal{V}_i^b$) define the positions of the apical vertices, which without a loss of generality we assume lie on the plane $\boldsymbol{x} \cdot \hat{\boldsymbol{z}} = 0$ (i.e. the $x - y$ plane). Here $\hat{\boldsymbol{z}}$ denotes the unit vector in the z axis.

$$\boldsymbol{x}_{\boldsymbol{q}}^a = \boldsymbol{x}_{\boldsymbol{q}}^b - (\boldsymbol{x}_{\boldsymbol{q}}^b \cdot \hat{\boldsymbol{z}})\hat{\boldsymbol{z}} \tag{12}$$

where the superscripts $a$ and $b$ denote apical and basal vertices respectively. Indeed, let the superscript $a$ be the above projection to the $xy$ plane of any basal position denoted with the superscript $b$.

Let $\mathcal{V}_i$ denote the vertex indices that are attributed to a given cell $i$, and $\mathcal{T}_q$ denote the set of cell indices that are attributed to a vertex $q$. (Note that the size of $\mathcal{T}_q$ is always 3, given in the model cells always meet at tricellular junctions).

###### A.1.1 Boundary conditions

We consider a tissue with $N$ cells, where the $x$ and $y$ dimensions are periodic, within a box of dimension $(L \times L)$. In the rest of the text, all distances are calculated respecting these periodic boundary conditions, which can be calculated via the periodic displacement (denoted with superscript $P$):

$$(\boldsymbol{x}_i - \boldsymbol{x}_j)^P = \boldsymbol{\Delta x}_{\boldsymbol{i,j}}^P = \quad \bmod\left(\boldsymbol{x}_i - \boldsymbol{x}_j + \frac{L}{2}, L\right) - \frac{L}{2} \tag{13}$$

$$\|\boldsymbol{x}_i - \boldsymbol{x}_j\|^P = \|\boldsymbol{\Delta x}_{\boldsymbol{i,j}}^P\| \tag{14}$$

We will drop the $P$ superscript from hereon.

###### A.1.2 Apical surfaces

**Apical perimeter**

The apical perimeter of cells – the subject of cortical tension – is defined by:

$$P_i^a = \sum_{q \in \mathcal{V}_i} \|\boldsymbol{x}_q^a - \boldsymbol{x}_{q+1}^a\| \tag{15}$$

where $(q + 1)$ is the counter-clockwise adjacent vertex to $q$ for cell $i$.

**Apical area**

Likewise the apical area is calculated using the shoelace theorem

$$A_i^a = \frac{1}{2} \sum_{q \in \mathcal{V}_i} (\boldsymbol{x}_q^a \times \boldsymbol{x}_{q+1}^a) \cdot \hat{\boldsymbol{z}} \tag{16}$$

While the reference frame is irrelevant for this calculation, for stability (given periodic distance calculations), we choose to calculate the areas with respect to an approximation of the apical centre of mass $\tilde{\boldsymbol{r}}_i^a$ (which is defined below), which is the voronoi centre of the polygon for the first iteration (see initialisation) or the value of $\boldsymbol{r}_i^a$ from the previous time step for all other iterations.

$$A_i^a = \frac{1}{2}\sum_{q\in\mathcal{V}_i}((\boldsymbol{x}_q^a - \tilde{\boldsymbol{r}}_i^a)^P \times (\boldsymbol{x}_{q+1}^a - \tilde{\boldsymbol{r}}_i^a)^P)\cdot\hat{\boldsymbol{z}} \tag{17}$$

**Apical centre of mass**

For later calculations, it is also important to calculate the apical centre of mass. This is calculated by the following general equation:

$$\boldsymbol{r}_i^a = \frac{1}{6A_i^a}\sum_{q\in\mathcal{V}_i}(\boldsymbol{x}_q^a + \boldsymbol{x}_{q+1}^a)\left[(\boldsymbol{x}_q^a \times \boldsymbol{x}_{q+1}^a)\cdot\hat{\boldsymbol{z}}\right] \tag{18}$$

which we adapt as an update equation (converging in a single iteration), to allow for stability given periodic distances

$$\boldsymbol{r}_i^a = \tilde{\boldsymbol{r}}_i^a + \frac{1}{6A_i^a}\sum_{q\in\mathcal{V}_i}((\boldsymbol{x}_q^a - \tilde{\boldsymbol{r}}_i^a)^P + (\boldsymbol{x}_{q+1}^a - \tilde{\boldsymbol{r}}_i^a)^P)\left[((\boldsymbol{x}_q^a - \tilde{\boldsymbol{r}}_i^a)^P \times (\boldsymbol{x}_{q+1}^a - \tilde{\boldsymbol{r}}_i^a)^P)\cdot\hat{\boldsymbol{z}}\right] \tag{19}$$

##### A.1.3 Basal surfaces

While apical surfaces are constrained to be coplanar by default, given basal surfaces will be typically comprised of more than three vertices, geometrically, they need not be coplanar. We therefore triangulate the basal surface with respect to a basal centroid $\boldsymbol{r}_i^b$.

**Basal centroid**

The basal centroid is a projection of the apical centre of mass $\boldsymbol{r}_i^a$ in the direction of $\hat{\boldsymbol{z}}$, where the value $r_{z,i}^b = \boldsymbol{r}_i^b \cdot \hat{\boldsymbol{z}}$ is calculated as the angular mean $z$ along the locus of points that comprise the basal perimeter. This enforces the basal centroid to exist as close as possible to the average plane of the basal vertices. Other energy terms in the model (see basal bending section) mean that the details of this calculation are unimportant, as basal vertices are encouraged to exist on a plane (which may take any orientation).

Let $\theta_{q,i}$ denote the signed angle of the apical vertex $q$ and the $x$ axis, with respect to the apical centre of mass i.e.

$$\theta_{q,i} = \text{atan2}\left((\boldsymbol{x}_q^a - \boldsymbol{r}_i^a)\cdot\hat{\boldsymbol{y}}, (\boldsymbol{x}_q^a - \boldsymbol{r}_i^a)\cdot\hat{\boldsymbol{x}}\right) \tag{20}$$

Where $-\pi \leq \theta_{q,i} \leq \pi$

Then let $z(\theta)$ be the $z$ component of the basal edge at a particular $\theta$ i.e.

$$z(\theta) = \boldsymbol{x}_q^b \cdot \hat{\boldsymbol{z}}\frac{(\theta - \theta_{q,i})(\boldsymbol{x}_{q+1}^b - \boldsymbol{x}_q^b)\cdot\hat{\boldsymbol{z}}}{\theta_{q+1,i} - \theta_{q,i}} \;\; ; \text{for } \theta_{q,i} < \theta \leq \theta_{q+1,i} \tag{21}$$

Then $r_{z,i}^b$ is defined as the angular mean:

$$r_{z,i}^b = \frac{1}{2\pi}\int_0^{2\pi} z(\theta)d\theta \tag{22}$$

This reduces to:

$$r_{z,i}^b = \frac{1}{4\pi}\sum_{q\in\mathcal{V}_i}(\theta_{q+1,i} - \theta_{q,i})\left((\boldsymbol{x}_{q+1}^b + \boldsymbol{x}_q^b)\cdot\boldsymbol{z}\right) \tag{23}$$

Finally, we can build $\boldsymbol{r}_i^b$ by:

$$\boldsymbol{r}_i^b = \boldsymbol{r}_i^a + r_{z,i}^a\hat{\boldsymbol{z}} \tag{24}$$

**Basal Curvature**

If basal vertices do not lie on a plane, the basal surface will be curved/bent. To quantify this, we approximate the curvature of the basal surface as $\phi_i^b$

$$\phi_i^b = \sum_{q\in\mathcal{V}_i}\arccos\left(\frac{(\boldsymbol{x}_q^b - \boldsymbol{r}_i^b)^P\cdot(\boldsymbol{x}_{q+1}^b - \boldsymbol{r}_i^b)^P}{\|\boldsymbol{x}_q^b - \boldsymbol{r}_i^b\|^P\|\boldsymbol{x}_{q+1}^b - \boldsymbol{r}_i^b\|^P}\right) \tag{25}$$

By construction, $\phi_i \geq 2\pi$, meaning the differential $\|\phi_i - 2\pi\|$ describes how much the basal surface is curved/bent. When all basal vertices are coplanar, $phi_i = 2\pi$.

###### A.1.4 Volume

Together, these geometrical constructs define the volume of the cell $V_i$. This is decomposed into a set of $n_{vertex,i} \times 3$ tetrahedrons:

$$V_i = \frac{1}{6} \sum_{q \in \mathcal{V}_i} \left[ ((\boldsymbol{x}_q^b - \boldsymbol{r}_i^b)^P \times (\boldsymbol{x}_{q+1}^b - \boldsymbol{r}_i^b)^P \cdot \hat{\boldsymbol{z}})((\boldsymbol{x}_{q+1}^b + \boldsymbol{x}_{q+1}^b) \cdot \hat{\boldsymbol{z}}) + r_{z,i}^b \|(\boldsymbol{x}_q^b - \boldsymbol{r}_i^b)^P \times (\boldsymbol{x}_{q+1}^b - \boldsymbol{r}_i^b)^P\| \right] \tag{26}$$

##### A.2 Energy functional and dynamics

To model the dynamics of the tissue, we presume that the vertices move according to overdamped dynamics, under an energy functional $\mathcal{E}$.

We define $\mathcal{E}$ to reflect the forces inherent to an epithelial tissue such as ours. We consider two energies: $\mathcal{E}_{internal}$, representing the internal forces in the tissue; and $\mathcal{E}_{external}$ to represent the external forces applied to the tissue. Here, $\mathcal{E} = \mathcal{E}_{internal} + \mathcal{E}_{external}$

###### A.2.1 Internal energy

$$\mathcal{E}_{internal} = \sum_i \kappa_V (V_i - V_{i,0})^2 + \kappa_A f(A_i^a)^2 + T_{cell,i}^a P_i^a + F_{bend}(\phi_i^b - 2\pi)^2 \tag{27}$$

To break this down:

- $\kappa_V (V_i - V_{i,0})^2$ : volume preservation with an optimal volume for a particular cell $i$ being $V_{i,0}$. $\kappa_V$ is the associated bulk modulus. We choose $V_{i,0} = 1 \forall i$ and set $\kappa_V$ to be higher than the other parameters, such that cell volume is as close as possible to 1 to align with the observation of incompressibility of cells, and volume-invariant shape changes.
- $\kappa_A$ : apical area elasticity/surface tension.
- $T_{cell,i}^a P_i^a$: line tension on apical perimeter, with force $T_{cell,i}^a$.
- $\alpha(\phi_i^b - 2\pi)^2$ bending energy for the basal surface. $\alpha$ is chosen to be high such that the basal surface approximates a flat plane. (Note that if we were to calculate $\phi_i^a$ it would equal $2\pi$ by construction).

###### A.2.2 External energy

Further, we presume an external force applied to the tissue, which takes the form of an isotropic stretch, in a similar manner to [Bielmeier et al., 2016]:

$$\mathcal{E}_{external} = T_{external} A^a \tag{28}$$

where $A^a = \sum_i A_i^a$.

###### A.2.3 Tissue growth

To model tissue growth/stretching given periodic boundary conditions, we follow [Guerrero et al., 2019, Bocanegra-Moreno et al., 2023] and consider the periodic box size $L$ as a dynamic variable that is also subject to change under the energy functional.

#### A.3 Vertex position evolution

In the true coordinate space, the evolution of the vertices are given by:

$$\partial_t \boldsymbol{x}_i = -\mu_x \frac{\partial E}{\partial \boldsymbol{x}_i} \tag{29}$$

Where $\boldsymbol{x}_i$ is a shorthand for $\boldsymbol{x}_i^b$, which is sufficient to define the entire geometry of the cell (see above)

Instead, we calculate the dynamics in the rescaled coordinate space, which effectively transforms the damping parameter $\mu_x$.

$\frac{\partial E}{\partial \hat{\boldsymbol{x}}_i}$ is solved using a reverse-mode auto-differentiation using the function *jax.jacrev* [Bradbury et al., 2018].

#### A.4 T1 transitions

While we anticipate T1 transitions to be infrequent in our simulations under appropriate regimes, including them is necessary such that vertices don't cross over each other. Given the 3D geometry of our tissue is achieved by planar projections, T1 transitions need only be solved from the apical domain. We follow standard practices to perform T1 transitions. Briefly:

- If an (apical) edge is smaller than a length $\epsilon_{swap}$ then a T1 transition is performed
- This is achieved by calculating the mid-point of the corresponding basal edge $\boldsymbol{x}_{mid} = (\boldsymbol{x}_q^b + \boldsymbol{x}_{q+1}^b)/2$, identifying the normal of the tangent line of the apical edge:($\boldsymbol{n} = (\boldsymbol{x}_q^a - \boldsymbol{x}_{q+1}^a)^P \times \hat{\boldsymbol{z}}$; $\hat{\boldsymbol{n}} = \boldsymbol{n}/\|\boldsymbol{n}\|$)
- Projecting the new vertices by a distance $\epsilon_{replace}$ along this vector (i.e. $\boldsymbol{x}_{mid} \pm \epsilon_{replace}\hat{\boldsymbol{n}}$. Here, $\epsilon_{replace}$ is chosen such that $2\epsilon_{replace} > \epsilon_{swap}$, meaning T1s don't immediately switch back in the next iteration.

### B Local and global contributions to cell stretch

To motivate this minimalistic theory, we examine the forces experienced by a cell embedded in a heterogeneous tissue to make the argument that Notch-mediated dampening of cell tension can have non-local and global effects, mechanically coupling cells and allowing them collectively to transition towards a stretching regime.

Considering a uniform hexagonal lattice of cells, we analyse the energy differential upon an isotropic stretch:

$$\boldsymbol{x_j} = \boldsymbol{x_j^0} + l\frac{(\boldsymbol{x_j^0} - \boldsymbol{r_i})}{\|(\boldsymbol{x_j^0} - \boldsymbol{r_i})\|} \tag{30}$$

where the superscript $s^0$ denotes the value in the base state. Under the definition $l = \hat{l}L$, this leads to a decomposition

$$\frac{\partial E}{\partial l} = \underbrace{L\frac{\partial E}{\partial \hat{l}}\bigg|_L}_{\text{Local rearrangement}} + \underbrace{\hat{l}\frac{\partial E}{\partial L}\bigg|_{\hat{l}}}_{\text{Global stretch}} \tag{31}$$

The first term considers the component of stretch that preserves the area of the tissue by locally moving vertices. The second term considers the component of stretch that changes the area of the tissue.

With isotropic stretching, we can calculate the changes in area and perimeter for the cell in question ($i$) and its neighbours ($k$), while preserving the global area of the tissue. Let $\epsilon = \frac{l}{l+d_{i\to j}}$, where we define $d_{i\to j} = \|(\boldsymbol{x_j^0} - \boldsymbol{r_i})\|$

$$A_i = A_i^0(1+\epsilon)^2 \text{ Area of cell in question} \tag{32}$$
$$P_i = P_i^0(1+\epsilon) \text{ Perimeter of cell in question} \tag{33}$$
$$A_k = A_k^0 - \frac{\sqrt{3}}{4}(\ell_{ik}^0)^2\epsilon(2+\epsilon) \text{ Area of neighbouring cell } k \tag{34}$$
$$P_k = P_k^0 - \ell_{ik}^0\epsilon \text{ Perimeter of neighbouring cell } k \tag{35}$$
$$\tag{36}$$

where $\ell_{ik}$ is the edge length of the two vertices cells $i$ and $k$ share.
Expanding the first term in Eqn. 31, describing local rearrangements, we get

$$\frac{\partial E}{\partial \hat{l}} = \frac{d_{i\to j}}{l + d_{i\to j}}\frac{\partial E}{\partial \epsilon} \tag{37}$$

where

$$\frac{\partial E}{\partial \epsilon} = \frac{\partial E_i}{\partial \epsilon} + \sum_{k\in\mathcal{N}_i}\frac{\partial E_k}{\partial \epsilon} \tag{38}$$
$$=2\kappa_P\left[(P_i - P_0)P_i^0 - \sum_{k\in\mathcal{N}_i}(P_k - P_0)\ell_{ik}^0\right] + \left[T_{cell_i}P_i^0 - \sum_{k\in\mathcal{N}_i}T_{cell,k}\ell_{ik}^0\right] + 2\kappa_A(1+\epsilon)\left[f'(A_i)A_i^0 - \sum_{k\in\mathcal{N}_i}\frac{(\ell_{ik}^0)^2\sqrt{3}}{4}f'(A_k)\right] \tag{39}$$

with $\mathcal{N}_i$ defining the neighbours of cell $i$, and

$$f'(A) = \begin{cases} A, \text{ if } A > A_0 \\ 2(A - A_0) \text{ otherwise}\end{cases} \tag{40}$$

Here, one can see how the mechanics of the neighbouring cells affects the capacity of a chosen cell to stretch. One can also observe that in a hexagonal lattice, where all cells have the same parameters, $A_i, P_i = 0\ \forall i \implies \frac{\partial E}{\partial \epsilon} = 0$ i.e. the homogeneous hexagonal lattice is in force-balance (noting that $\sum_k \ell_{ik}^0 = P_i^0$ and $\sum_k \frac{(\ell_{ik}^0)^2\sqrt{3}}{4} = A_i^0$
Additionally, we can expand the second term, considering global stretch:

$$\frac{\partial E}{\partial L} = \sum_i \frac{\partial E_i}{\partial L} \tag{41}$$
$$= \sum_i \left[p_i\left(2\kappa_P(p_iL - P_0) + T_{cell,i}\right) + 2a_iL(\kappa_A f'(a_iL^2) - T_{external})\right] \tag{42}$$

defining the dimensionless areas and perimeters as $a_i = A_iL^2; p_i = P_iL$. Therefore, cell stretch has contributions from the energies of all cells, and will be influenced by the frequency of Notch via changes in the average $T_{cell,i}$.
